# Chemically defined lipid diets reveal the versatility of lipidome remodeling in genomically minimal cells

**DOI:** 10.1101/2024.10.04.616688

**Authors:** Nataliya Safronova, Lisa Junghans, Jana Oertel, Karim Fahmy, James P. Saenz

## Abstract

All cells are encapsulated in a lipid membrane that provides a responsive interface between life and its environment. Although simple membranes can be built from a single type of lipid, cellular membranes contain 10s to 100s of unique lipid species. Deciphering the significance of lipidome complexity is a central challenge in understanding the design principles of living membranes. While functions of individual lipids have been extensively studied, understanding how lipidomes collectively contribute to membrane function and cell phenotypes is experimentally challenging in most organisms. To address this challenge, we turned to the simple pathogenic organism *Mycoplasma mycoides* and its genomically derived “Minimal Cell” JCVI-syn3B, to establish a living minimal membrane model system in which lipidome complexity can be experimentally manipulated. By complexing lipids with cyclodextrins, we introduce a chemically defined approach to deliver lipid ‘diets’ with different chemistries to cells, resulting in cellular lipidomes with as few as seven to nearly 30 lipids species. We explored how lipidome size and composition influences cell growth, osmotic sensitivity, and membrane adaptability to changes in growth temperature. Our findings indicate that lipidome composition dictates membrane adaptation to temperature change. Moreover, we show that lipidome diversity enhances cellular robustness to hypoosmotic shock. We further show that impaired acyl chain remodeling in the minimal cell is associated with impaired membrane temperature adaptation. Finally, we demonstrate as a proof of principle, how cells with tuneable lipidomes can be used as experimental chassis for screening membrane active antimicrobial peptides. Our study introduces an experimental resource and foundation for deciphering the role of lipidome complexity in membrane function and cellular fitness.

## Introduction

Lipid membranes serve as responsive interfaces to compartmentalize life and as platforms for cellular bioactivity. The unique material properties of lipid bilayers allow cells to build a surface membrane that is both mechanically stable to withstand environmental perturbations, and fluid enough to support bioactivity. The simplest membrane can be assembled from only one type of lipid, but biological membranes are built from much wider arrays of lipid species^1,2^. Why have cells evolved to make membranes that are so complex? Complexity is one of the features of biological systems that contributes to robustness^3^. Thus, the complexity of cellular lipidomes may have evolved to enhance robustness of the membrane to perturbations or noise. More complex lipidomes also have a higher degree of freedom to modulate their properties and function, since each variation in lipid structure can uniquely alter the physical state of the bilayer. More degrees of freedom could give the cell flexibility to independently optimize different membrane properties such as permeability, fluidity, or thickness. While much is known about how structures of individual lipids contribute to bilayer properties, how lipidome complexity collectively determines membrane robustness and physical homeostasis is an open question of central importance to the design principles of living membranes.

The lipid membrane and its broad physical and chemical features are a conserved feature of all cells, suggesting that it was present in the last universal common ancestor, and possibly early in the emergence of life^4^. Elucidating the basis for lipidome complexity and the role of lipids in supporting life requires bridging a gap between the staggeringly complex and experimentally challenging cellular lipidomes of modern organisms^5,6^, and simpler lipid mixtures that may have supported ancient life.

While functions of individual lipids are widely studied, understanding the collective role of lipidome complexity in living organisms is experimentally and conceptually challenging. For example, the genetic complexity of even relatively simple prokaryotic membrane research models, such as *B. subtilis*^7^, *E. coli*^8^ and *M. extorquens* poses considerable challenges for large-scale lipidome manipulations, which often result in the accumulation of lipid biosynthetic precursors^9^, non-linear metabolic feedbacks^10,11^, and/or other genetic artefacts. Further, the presence of intracellular membrane compartments in both prokaryotic^12^ and eukaryotic^13^ cells complicates experimental isolation, perturbation and biophysical analysis of an individual membrane. To address these limitations inherent to common research model organisms, we aimed to develop a minimal cellular model system with a defined, simple, yet compositionally flexible and living membrane. As we will show, the Minimal Cell (JCVI-syn3B) and its pathogenic parent organism *Mycoplasma mycoides* comprise a robust system that is uniquely poised to deciphering the role of lipidome complexity in the design principles of living membranes.

The isolation and characterization of Mycoplasmas^14,15,16^ revealed the extreme evolutionary reduction of their biosynthetic capacity, from amino acids to lipids. The remarkable dependence of these pathogens on the host nutrient supply, including lipids, provides an unprecedented opportunity to control their lipidome composition through externally supplemented lipids, thereby obviating the need for genetic manipulation. Several decades ago, pioneers of Mycoplasma research at the Hebrew University conducted a tremendous effort in examining Mycoplasmas and establishing their potential as the simplest living membrane models^16,17,18^. However, comprehensive control of Mycoplasma lipid composition has not yet been fully realized and the advantages of a tunable minimal living membrane model system have not yet been fully explored. The synthetic reduction of *Mycoplasma mycoides* genome to the minimal gene set led to the creation of the minimal cell, JCVI-syn3B. Our lipidomic insights suggest impaired mechanisms of lipidome homeostasis in the minimal cell; nevertheless, this organism is a self-replicating organism, capable of withstanding environmental perturbations and extensive lipidomic remodeling^24^. Syn3B retains the experimentally advantageous features of Mycoplasmas, while providing insight into the fundamental requirements for minimal life. We have constructed a minimal membrane model system by utilizing JCVI-syn3B and its parental organism *Mycoplasma mycoides* to study the role of lipidome size and complexity in membrane adaptation and cellular growth.

## Results

### 1. Mycoplasma as a model system with controlled membrane lipid composition

Cellular membranes employ a wide range of structurally distinct lipid species, ranging from 10s to 100s of lipids, generating a large variability in complexity of membrane lipidomes, both in terms of size and composition^6,2,1^. Compositionally, lipids can be divided into major classes (e.g. sterols, phospholipids and sphingolipids and further sub-classes, e.g. phospholipids with different head groups (Fig.1a). Each lipid class is characterized with class-specific structural features, such as acyl chain length and degree of unsaturation in phospholipids (Fig.1a). It is useful to consider acyl chain features separately by class, as the properties of lipids are dependent on the combinatorial specificity of both head groups and acyl chains^1,19^.

**Figure 1.**
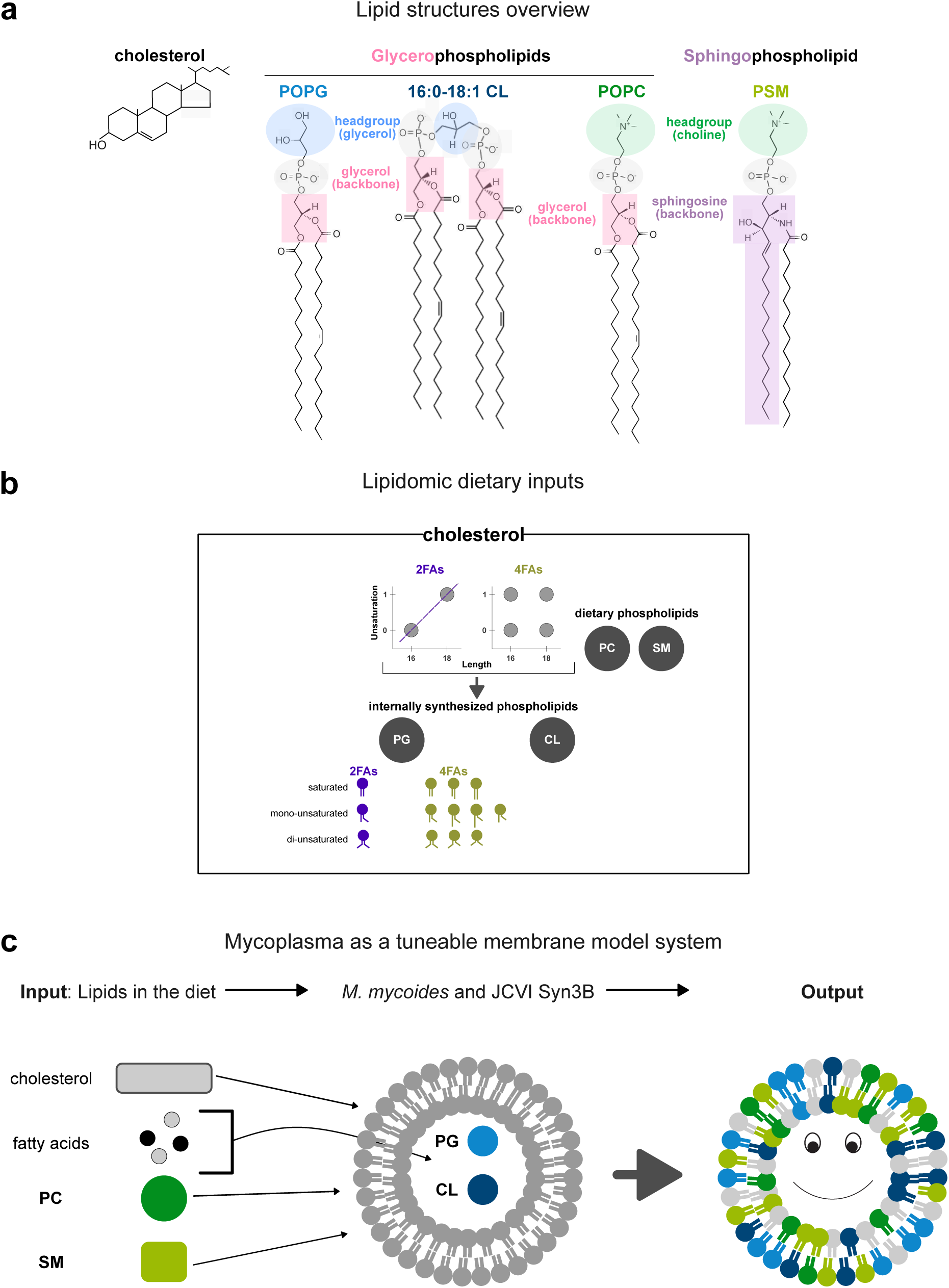
Key aspects of defined lipid diet methodology. **a**. Chemical structures of lipid classes, represented in this study. Top: cholesterol. Bottom: POPG (palmitoyl-oleoyl-phosphatidylglycerol), 16:0-18:1-16:0-18:1 cardiolipin, POPC (palmitoyl-oleoyl-phosphatidylcholine), PSM (phosphatidylsphingomyelin). **b**. The overview of lipids, fed to mycoplasma in this work. Cholesterol is delivered at constant concentration to all lipid diets. The minimal diet - 2FAs - contains palmitic acid (C16:0) and oleic acid (C18:1). 4FAs diet delivers palmitoleic acid (C16:1) and stearic acid (C18:0). These fatty acids allow mycoplasma to produce internal PG and CL species. In addition, POPC and egg SM were added to fatty acids to expand mycoplasma lipid repertoire.

To establish a minimal living membrane model system in which lipidome size can be experimentally manipulated, we took advantage of the limited capacity for *M. mycoides* and *Syn3.0B* to synthesize their own lipids, and their resulting dependence on exogenous lipid uptake^17^. Mycoplasma cannot synthesize fatty acids^16^. However, when free fatty acids are provided in the growth media, Mycoplasma can synthesize phosphatidylglycerol (PG) and cardiolipin (CL)^20^ (Fig.1c). *M. mycoides* and Syn3B also require a sterol (e.g. cholesterol) for growth, which must be delivered exogenously^21,22^. Thus, the minimum lipid repertoire of *M. mycoides* or Syn3B is cholesterol, PG and CL. Additionally, *M. mycoides* and Syn3B can take up phospholipids and sphingolipids that are provided in the media^23,24^. Using methyl-beta cyclodextrins to complex fatty acids and cholesterol and methyl-alpha cyclodextrins to complex phospholipids we delivered chemically defined mixtures of lipids to *M. mycoides* and JCVI-Syn3B. Thus, by creating defined “lipid diets” with varying fatty acid or lipid classes, we manipulate the lipidome complexity of *M. mycoides* and Syn3B.

The minimal lipid diet that can support growth of *M. mycoides* is cholesterol, and two free fatty acids: palmitate and oleate (C16:0 and C18:1)^25,26^. We used this lipid diet, designated as ‘2FAs’, as the minimal chassis to build upon. A fundamental constraint of this diet is that variations in double bonds and chain length are coupled, thereby limiting the lipidomic degrees of freedom (Fig. 1b). To examine the effect of this constraint on lipidome flexibility we designed a lipid diet with four fatty acids (4FA: C16:0, C16:1, C18:0, C18:1) which expands the range of acyl chain combinations available for each of the cell-synthesized phospholipid classes (Fig. 1b) and allows independent variation of double bonds and chain length (Fig. 1b).

The complexity of lipid classes can further be controlled through lipid diet. When grown on fetal bovine serum (FBS) as an undefined complex lipid diet, *M. mycoides* will incorporate exogenous phosphatidylcholine and sphingomyelin into their lipidomes^24^. Therefore, to examine the effect of class complexity we designed three lipid diets based on the 2FA diet containing additionally phosphatidylcholine (2FA-PC), sphingomyelin (2FA-SM), or both PC and SM with either 2FAs (2FA-PC-SM) or 4FAs (4FAs-PC-SM). For PC we used a single species of PC with one palmitoyl (C16:0) and one oleoyl (C18:1) acyl chain (POPC: 1-palmitoyl-2-oleoyl-sn-glycero-3-phosphocholine, structure shown in Fig.1a. For SM diets we used egg sphingomyelin, which is a mixture of N-hexadecanoyl-D-erythro-sphingosylphosphorylcholines composed of >90% of one SM species with a C16:0 acyl chain (chemical structure shown on Fig.1a) (the precise species composition of egg SM is shown on Supplementary Figure 2b).

### 2. Lipid diets shape *M. mycoides* and JCVI-syn3B cell size and growth rate at 37°C

To evaluate the effect of varying lipid complexity on the growth of *M. mycoides* and Syn3B, we recorded heat flow curves through microcalorimetry at 37°C (SFig. 1b) and calculated the respective growth rates to quantitatively assess how varying lipid diets influenced *M. mycoides* and Syn3B growth (Fig.2a). On average, *M. mycoides* shows faster growth rate on all diets, than its synthetic counterpart; remarkably, diets with only fatty acids and cholesterol have the opposite effect on *M. mycoides* and Syn3B growth, with *M. mycoides* displaying the fastest growth on the minimal diet. At the same time, diets with sphingomyelin yield comparable growth in *M. mycoides*, while Syn3B displays the best growth with 2FAs-PC-SM diet. In contrast to *M. mycoides*, Syn3B grows faster in presence of external phospholipids in the diet, which might be the result of lower energy demands associated with uptake rather than synthesis of phospholipids. Average differences in growth rates, produced by the same lipid diet in two organisms, suggest that the vast reduction in genome size of the minimal cell is a crucial factor in shaping bacterial growth, which agrees with growth rates dynamics of mycoplasma cells, grown on undefined lipid diet (FBS)^24^. However, for some diets (2FAs-PC, 2FAs-PC-SM) these variations are negligible, in contrast to FBS results. In addition, variations in lipid diets yield 2-3-fold difference across lipid diets; taken together, these observations showcase the fundamental importance of lipid repertoire for the cellular fitness.

**Figure 2.**
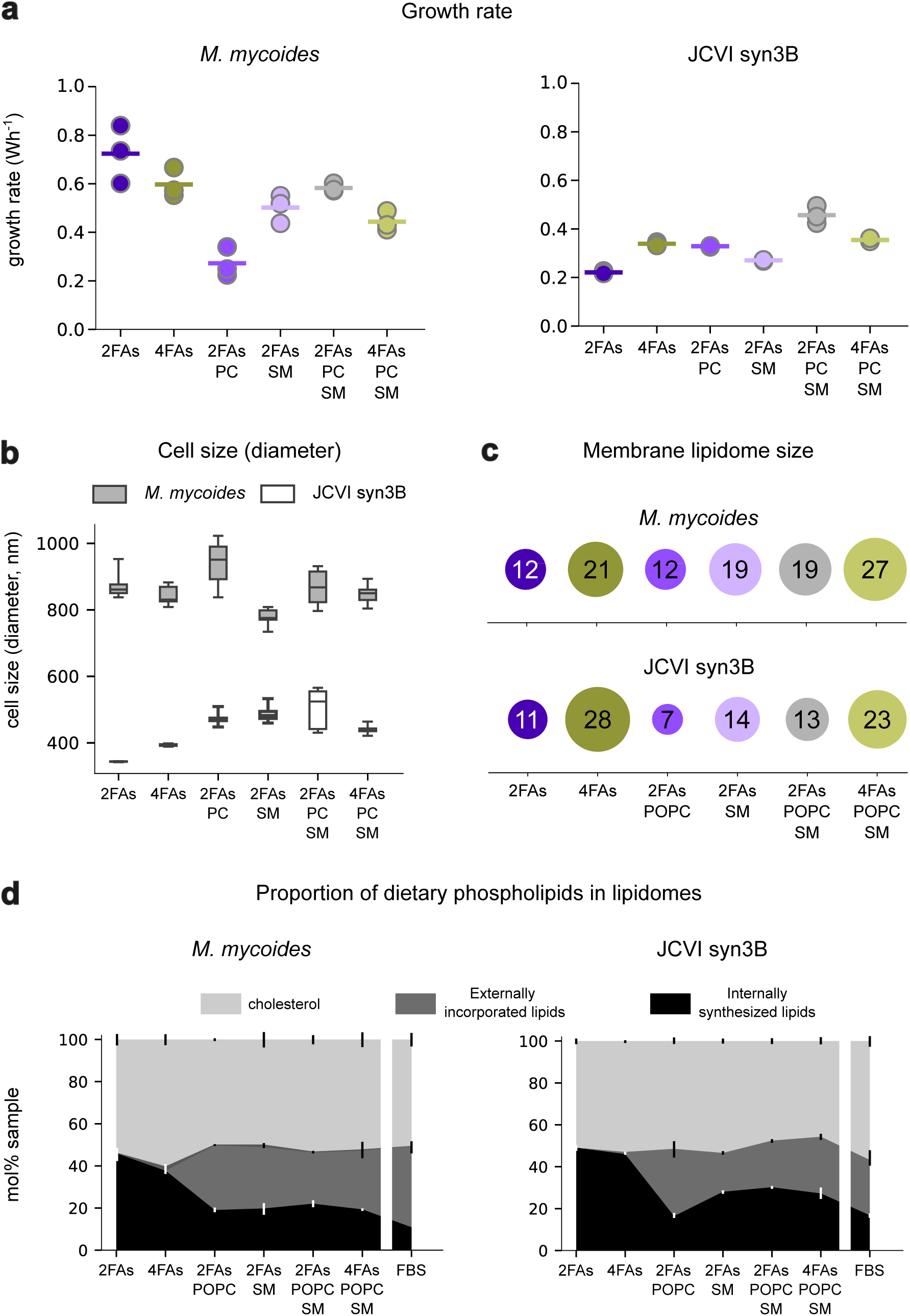
Lipid diet defines growth rate, cell size and membrane lipidome composition. **a.** *M. mycoide*s and Syn3B growth rate at 37°C, n = 3**. b.** Mycoplasma cell size (diameter) in nm. n = 9. **c**. Mycoplasma lipidome size on each of the defined lipid diets, represented as the number of structurally distinct lipid species in each lipidome. **d**.Proportion of cholesterol (top), dietary external phospholipids (middle) and internally synthesized lipid lipids (bottom) in each of the lipid diets. n = 3, standard deviations shown as bars. Lipidome of cells grown on FBS, grouped by cholesterol, external and internal phospholipids, is shown for reference.

To further emphasize the importance of lipid repertoire in cellular metabolism, we recorded the average cell size using dynamic light scattering (Fig.2b). It is established that bacterial growth rate is closely coupled with cellular volume^27,28^, and, at the same time, cell size is coupled with lipid synthesis^29^. Moreover, previous experiments on a non-pathogenic relative of mycoplasma, *Acholeplasma laidlawii*, showed that supplementing the bacterium with cholesterol or unsaturated fatty acids yielded larger cell size^30^. Bacterial cells quickly adapt to changes in nutrient conditions by careful timing of cell division^31,32^, which keeps average cell size independent of the bacterial growth rate^31^. Our findings indicate the average cells size (diameter) of *M. mycoides* fluctuates around 800-900 nm and is conserved across lipid input variations. The reported cell diameters of different mycoplasma species vary from 0.15-1 um range^33^, which is consistent with our observations. *M. mycoides* does not show any correlation between average growth rate and cells size (SFig.1c). As expected, the minimal cell on average features about 2-fold lower cell size of around 400-450 nm, which agrees well with previous reports on the minimal cell parameters^34^. In contrast to *M. mycoides*, Syn3B features a distinct correlation between lipid input and cell size (SFig.1c). The minimal diet (2FAs) yields the smallest cells size of about 350 nm, which increases with more fatty acids, present in the diet (4FAs), consistent with observations in Acholeplasma cells^30^. Once a PC and/or SM are present in the diet, cell size increases further to an average of 420-450 nm. There is a non-linear positive correlation between the cell size and the growth rate in the minimal cell (SFig. 1c). The addition of PC and/or SM might enhance metabolic rates by redirecting energy from internal lipid biosynthesis; additionally, PC and SM can relieve membrane tension and curvature, associated with internally synthesized CL^35^. It therefore appears that the minimized genome in Syn3B cannot effectively regulate cell size and is more sensitive to changes in lipid input.

Having observed how varying lipid input influences mycoplasma cell size and growth rate at 37°C, our next step is to dissect mycoplasma lipidomes to show that lipid repertoire defines membrane lipid size and composition. Variations in lipid diet composition define the respective lipidome size of *M. mycoides* and JCVI-syn3B membranes (Fig.2c). Lipidome sizes range from as few as 7 lipid species (Syn3B, (2FAs-PC) diet) to nearly 30 lipid species in cells, grown on diets with 4FAs. The largest lipidomes, generated from 4FAs-containing lipid diets, approach the recently described lipidomic complexity of *M. extorquens*, which is still one of the smallest cellular lipidomes reported to date^1^. At the same time, defined lipid diets produce up to an order of magnitude smaller lipidome sizes, than cells grown with fetal bovine seum (FBS, an undefined complex lipid diet), demonstrating the wide potential range for tuning lipidome size. In both *M. mycoides* and Syn3B, the 2FAs-PC lipid diet produces the smallest lipidome size, which is nevertheless larger, than the smallest mycoplasma lipidome, ever reported^36^. The main reason behind the difference lies in the presence of free fatty acids in the defined lipid diets, which support mycoplasma’s endogenous lipid synthesis of PG and cardiolipin.

To examine the distribution of exogenously-derived lipids with varying lipid diet, we aggregated the lipidome in terms of externally-derived cholesterol and phospholipids (PC and SM) in endogenously synthesized glycerophospholipids (e.g. PG and cardiolipin)(Fig. 2d). By visualizing the lipidome in this simplified way, we highlight two general features of mycoplasma lipidomes: first, average cholesterol abundance is highly conserved at around 50-55 mol% of total lipidomes in all lipid diets. This value is in perfect alignment with sterol content of cells, grown on FBS^24^, showcasing cholesterol as a crucial structural component of *M. mycoides* and Syn3B membranes, regardless the lipidome size and composition. High cholesterol abundances in mycoplasma may compensate for the absence of a cell wall to support a robust barrier^37,38,39^. In addition, sterols have been implicated in the regulation of mycoplasmas membrane fluidity^37,40,41^. Second, both *M. mycoides* and Syn3B substitute about 20-30 mol% of their internally synthesized phospholipids with dietary PC and/or SM, lowering the PG and CL abundance to ∼20 mol% of total lipidome. This trend shows the strong preference of *M. mycoides* and Syn3B towards constructing their lipidomes from externally provided phospholipids. Nevertheless, the presence of both PG and CL in lipidomes under all conditions suggests that these lipids serve critical roles even at lower abundances.

### 3. Lipidome size and composition can be controlled through defined lipid diets

To dissect structural complexity of the Mycoplasma lipidomes, we parsed the lipidome in terms of lipid structural features, starting with lipid classes, followed by the phospholipid acyl chain-level features. Fig.3a shows a detailed lipid class composition of mycoplasma lipidomes. Changes in dietary lipid input can drive lipid class variations that follow similar trends in both *M. mycoides* and Syn3B (Fig.3a). In the absence of phospholipids from the lipid diet, *M. mycoides* synthesizes roughly equal amounts of PG and CL (20-25 mol%), while Syn3B prefers to incorporate more PG (30-40 mol% sample). External phospholipid abundances in different diets are relatively conserved in *M. mycoides* and Syn3B. Cells, grown on the most complex diets – 2/4FAs-PC-SM – exhibit a lipid class distribution, similar, yet not identical, to the lipidomic profiles of cells grown on undefined complex lipid source (FBS)^24^. In particular, the proportion of internally synthesized lipids (PG and CL) in both organisms remains higher with defined lipid input, and Syn3B tends to incorporate more PC. These results demonstrate how *M. mycoides* and Syn3B can tolerate major changes in lipid class composition, ranging from predominantly negatively charged bacterial lipid classes (PG, CL) to the zwitterionic PC and SM, characteristic of eukaryotic lipidomes.

**Figure 3.**
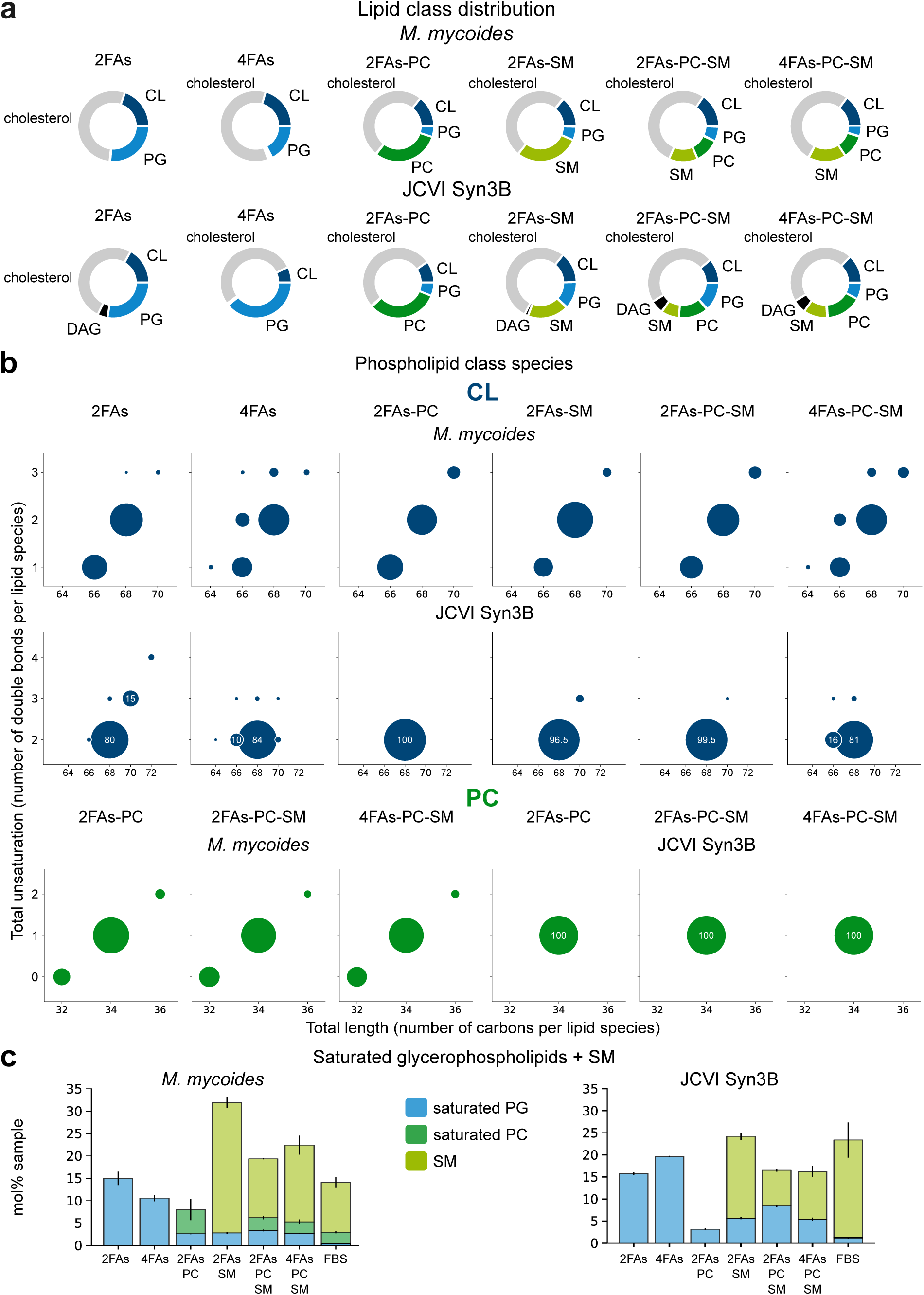
Mycoplasma membrane lipid composition at 37°C. **a.** Membrane lipid classes on each of the lipid diets in M. mycoides (top) and JCVI Syn3B (bottom). Pie charts illustrate an average mol% distribution of lipid classes in samples. n = 3. **b**. Cardiolipin (top) and POPC (bottom) species distribution in *M. mycoides* and Syn3B, plotted in acyl chain length-acyl chain unsaturation axes. Each bubble size shows an average mol% abundance of species in a class. n= 3. The detailed species distirbution for all classes can be found on Supplementary Figure xX. **c**. Saturated lipidome fractions in samples, mol%. Fractions are split in contributing phospholipids classes. mean +/- SD, n = 3.

Mapping phospholipidome species further illustrates how *M. mycoides* and Syn3B membrane lipid composition can be tuned through variations in lipid diet. The number of structurally distinct species within each lipid class is defined by the number of free fatty acid species provided through the diet. Larger fatty acid diversity in 4FAs diet expands the number of PG (SFig.2a) and CL (Fig.3b) species in both *M. mycoides* and JCVI-syn3B: while with 2FAs diets, the cells can only produce 3 PG species (SFig.2a), 4FAs diet supports the synthesis of 6-7 PG species, with similar patterns in CL acyl chain distribution. Fig.3b also shows that restricting lipid diet to only two fatty acid species prevents *M. mycoides* and Syn3B from independent regulation of acyl chain parameters in PG and CL. All of the internally produced PG and CL species with 2FAs diets fall on a straight line, implying a strict coupling between acyl chain length and unsaturation. In contrast, PG and CL species, synthesized in 4FAs and (4FAs-PC-SM) diets, arrange in a grid in length-unsaturation coordinates, breaking a strict linear codependence. Interestingly, while PG species profile is largely identical between *M. mycoides* and Syn3B (SFig.2a), the diversity of CL species in Syn3B tends to be more restricted and predominately consists of CL 68:2; in (2FAs-PC) diet, it is the only CL species (Fig.3b). In the rest of the diets, CL 68:2 comprises at least 80% of the total lipid class, even in the presence of 4FAs. This difference hints at the constraints on lipid synthesis and remodeling in the minimal cell, imposed by its minimized genome.

Furthermore, Syn3B lipidomes fed on 2FAs-POPC contain only POPC, while *M. mycoides* is capable of remodeling the acyl chains of POPC to generate more diverse acyl chain configurations resulting in several PC species (Fig.3b). Together with differences in CL species distribution, these results demonstrate the impaired Syn3B’s ability for lipidomic remodeling. The SM species distribution in both the lipid diet (egg SM) and the lipidomes of *M. mycoides* and Syn3B are comparable (SFig.2b), showing that there is no apparent remodeling or species-selective uptake of SM under the conditions considered in this study.

These results demonstrate that chemical properties and lipidomic complexity of *M. mycoides* and Syn3B membranes can be tuned using defined lipid diets varying in acyl chain or lipid class composition. We can further illustrate the flexibility of mycoplasma membrane model system by plotting the sum of saturated lipid species in all lipidomes (Fig.3c). The largest internally synthesized fraction of saturated, low-melting temperature lipids (represented by PG species only) is incorporated into lipidomes without external phospholipids in the diet and 2-3 folds lower in the lipidomes with POPC and/or SM. Such a difference might illustrate the need of mycoplasma to balance large quantities of low-melting temperature CL species in 2FAs and 4FAs diets. Egg SM in the diet contributes at least 10-15mol% abundance; as a result, mycoplasma tolerates the fluctuations in saturated lipidome fraction from about 5-10 mol% sample (2FAs-PC diet) to almost 35 mol% (2FAs-SM) diet. Large cholesterol abundance of at least half of total lipidome is likely to be one of the factors that allow mycoplasma such flexibility in constructing lipidomes with different chemical properties. Therefore, *M. mycoides* and Syn3B provide a range of minimal living membrane systems to explore physiological roles of lipidomic diversity and complexity.

### 4. Homeoviscous adaptation in mycoplasma is lipid-diet dependent

A core concept in membrane biology is homeoviscous adaptation, wherein membrane viscosity is conserved with varying temperature through lipidome remodeling^42^. In principle, such adaptation could be achieved with two lipid species, for example by varying the abundance of lipids with one and two double bonds. However, biomembranes exhibit much more complex patterns of temperature adaptation, involving tens of lipid species and class-specific acyl-chain remodeling^1,2,43,44^. Having established a method for tuning lipidome complexity through defined lipid diets, we examined their effects on homeoviscous adaptation (Fig.4a, b). To assay homeoviscous adaptation, we labelled mycoplasma membranes with and measured general polarization of Pro12A, a recently developed derivative of solvatochromatic dye c-laurdan^45^. The emission spectra of Pro12A is sensitive to the hydration of the bilayer, which in turn is tightly coupled with lipid order and membrane fluidity. By taking the ratio of two emission maxima, a general polarization (GP) index can be calculated, in which higher values correspond to lower fluidity. An advantage of Pro12A over c-laurdan is that because of its bulky polar head group, it selectively labels the outer leaflet of the plasma membrane, thereby preventing labeling of non-membranous hydrophobic compartments within the cell^45^. Pro12A therefore makes it possible to selectively monitor membrane fluidity in situ through bulk spectroscopic measurements, which is essential for small mycoplasma cells that challenge the resolution limits of microscopic analysis. The lipid order of *M. mycoides* and Syn3B membranes at 37°c and 30°C reported as Pro12A GP values are summarized on Fig.4a. On average, Pro12A GP of live mycoplasma cells are in the range of ∼0.25-0.45, which is a relatively high lipid order (low fluidity) and comparable with GP values for mammalian plasma membranes^45^ and agree perfectly with Pro12A GP values for mycoplasma cells, grown on FBS^24^. Syn3B membranes are more ordered across the temperature range, than *M. mycoides*, which is consistent with *M. mycoides* and Syn3B differences on undefined lipid diet.

**Figure 4.**
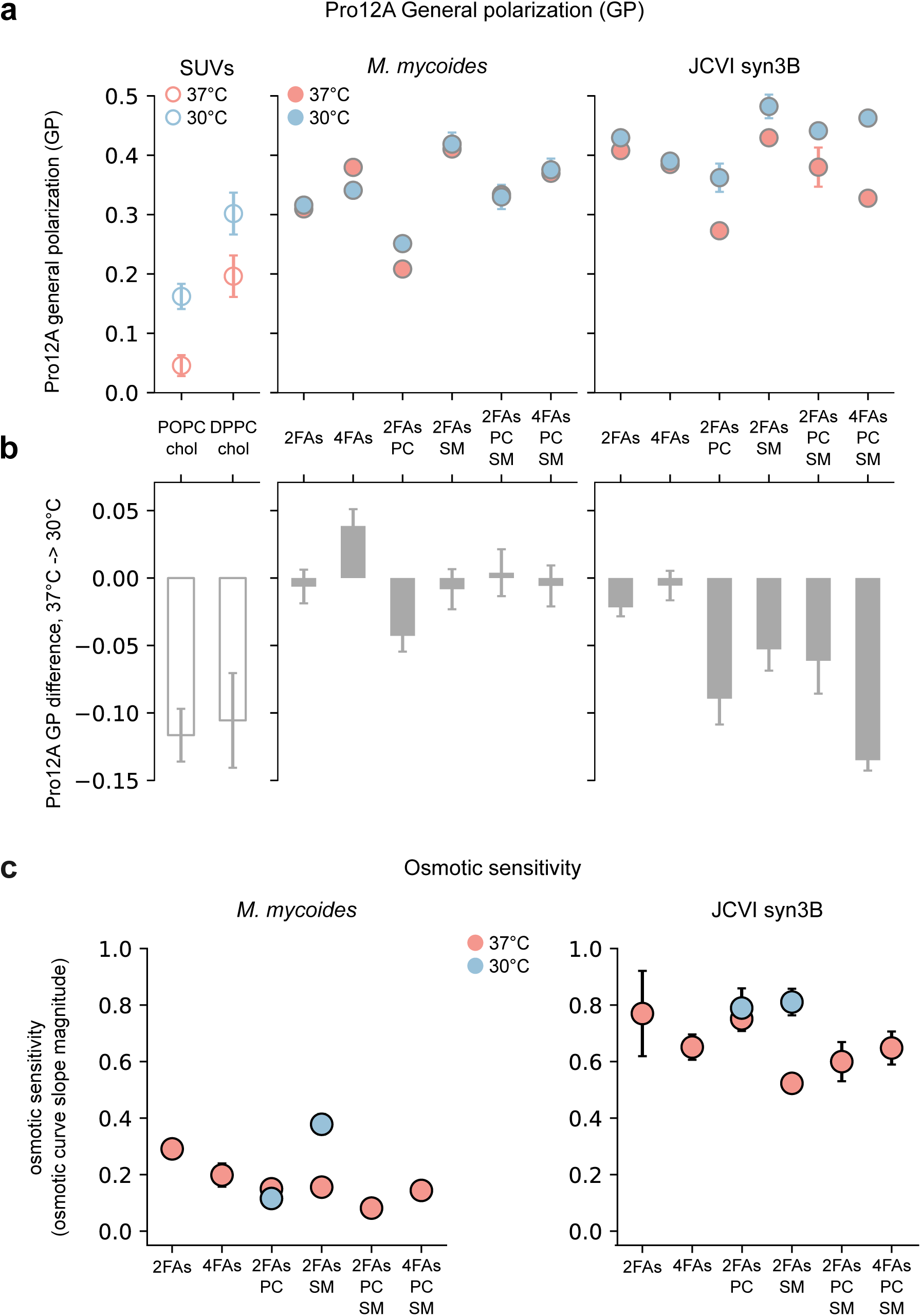
Lipid diet regulates homeoviscous adaptation and mechanical robustness of mycoplasma membranes. **a.** Pro12A GP of live mycoplasma membranes at 37°C and 30°C. The respective GP values of synthetic liposomes are shown for reference. mean +/- SD, n = 3. **b**. Pro12A GP difference, (37°C-30°C), calculated for synthetic membranes (left), *M. mycoides* (center) and Syn3B bars show mean difference +/- SD. **c**. Osmotic senstivity of live mycoplasma cells, grown at 37°C and 30°C.n = 3. The readout shows the absolute slope of salt-propidium iodide fluorescence curves. The curves are plotted on Supplementary Figure Xx.

The capacity for homeoviscous adaptation can be assayed by measuring the change in lipid order across different growth temperatures. We would expect lipid order (GP values) to remain constant if homeoviscous adaption were active. In the absence of homeoviscous adaptation, lipid order would increase with decreasing temperatures, and approach the change observed for non-adaptive liposomes, which in this case we have reconstituted from pure phospholipid and cholesterol. We plot the change in lipid order as the delta-GP for a given temperature range in Fig.4b. The change in lipid order for 37°C- 30°C temperature difference for *M. mycoides* is at least about 3-fold lower, than that for the synthetic liposomes GP, which is an indication that it effectively maintains membrane lipid order in a narrow range, when compared to non-adaptive membranes. The two bacterial strains show differences in their adaptation to temperature: in Syn3B, homeoviscous adaptation is surprisingly conserved in the absence of external phospholipids. The addition of PC and/or SM results in much lower membrane fluidity of Syn3B membranes at 30°C. In comparison to synthetic liposomes GP, 2FAs-PC and 4FAs-PC-SM diets provide almost no membrane fluidity regulation at lower temperature. How will such changes in membrane viscosity influence cellular robustness?

Membranes are important in environmental stress resistance^46^ and perturbed lipid biosynthesis can render cells more susceptible to physicochemical stresses, especially those directly targeting the membrane^47,48,49^. Osmotic shock directly challenges membrane rupture strength, and variations in osmotic susceptibility can indicate changes in membrane robustness. We therefore measured the sensitivity of *M. mycoides* and Syn3B grown on varying lipid diets at 37°C and 30°C (Fig.4c) to hypo-osmotic shock. To assess osmotic sensitivity of mycoplasma membranes, we subjected *M. mycoides* and Syn3B cells to varying buffer salinity, ranging from 200 mM NaCl (=isotonic) to 0 mM and used propidium iodide (PI) fluorescence as a proxy for the membrane rupture. Propidium iodide is a membrane-impermeable intercalating DNA stain, and is a useful indicator of disrupted membrane integrity of living cells^50^. We plotted the logarithmic PI’s fluorescence as a function of sample salinity (SFig. 3) and calculated the osmotic sensitivity as the linear slope of resulting curves. Fig.4c shows the absolute value of salt-fluorescence curves’ slope. Independent from the lipid input, *M. mycoides* displays much stronger robustness to hypoosmotic shock, than Syn3B (Fig.4c). While the minimal cell does not show a strong correlation between the lipid input and the respective osmotic sensitivity at 37°C, the addition of extra acyl chains in the diet of *M. mycoides* (2FAs-> 4FAs) leads to a decrease in osmotic sensitivity of about 30%. The presence of external phospholipids in the diet further reduces the sensitivity by 50-70%, compared to 2FAs diet. Therefore, the capacity to withstand the tensile stress, induced by the membrane swelling in hypoosmotic shock increases with larger structural degrees of freedom in lipid constituents,-either acyl chains (4FAs), or phospholipid headgroups (diets with PC, SM) or both (4FAs-PC-SM). In synthetic membranes, larger structural heterogeneity of lipid acyl chains increases bilayer compressibility and bending^51^, thus changing its responses to external stresses. At the same time, zwitterionic lipid bilayers, produced by mycoplasma in presence of PC and/or SM have different lipid-lipid and lipid-protein interaction dynamics^52,53^, than negatively charged membranes of *M. mycoides*, only producing PG and CL. Experiments with *S. cerevisiae* revealed that sphingomyelin enhances membrane resistance to osmotic shock through a variety of mechanisms, including cholesterol-SM microdomain modulation and calcium-channel activation. Taken together, our observations suggest that structural diversity of membrane lipids in *M. mycoides* enhances bacterial robustness in environmental perturbations, which is not reflected in the growth rate at 37°C (Fig.2a). Fig.4c therefore also exemplifies how a combination of readouts can be required to reveal the role of membrane lipidomic diversity in cellular fitness.

Varying the lipid input in mycoplasma not only modulates the capacity for homeoviscous adaptation, but also defines a wide range of membrane viscosity values at both 37°C and 30°C. 2FAs-PC diet results in the highest membrane fluidity (lowest Pro12A general polarization) at both temperatures and is less effective in supporting homeoviscous adaptation in *M. mycoides*. At the same time, 2FAs-SM diet yields the lowest membrane fluidity at both temperatures (the highest Pro12A general polarization value) in *M. mycoides* and Syn3B. There is nearly a two-fold average GP difference between 2FAs-PC and 2FAs-SM diets, showcasing how varying lipid input is capable of constraining membrane biophysical properties in mycoplasma membranes and in turn challenge its mechanical robustness at lower temperature. To demonstrate this, we additionally examined osmotic sensitivity of *M. mycoides* and JCVI-syn3B cells, grown with 2FAs-PC and 2FAs-SM diets at 30°C. Fig.4c shows that the higher membrane viscosity of 2FAs-SM cells results in large osmotic sensitivity. On the contrary, osmotic response at 30°C of cells, grown on 2FAs-PC diet, which remains unchanged at lower temperature. These observations demonstrate the key role of membrane lipid composition in shaping membrane robustness to environmental perturbations, such as lower growth temperature and osmotic shock. How does mycoplasma utilize its scarce lipidomic constituents on defined lipid inputs to adapt its membranes to lower growth temperature?

### 5. Patterns of lipidomic remodeling in mycoplasma

#### Lipid class variability

We have demonstrated that both *M. mycoides* and JCVI-syn3B employ remodeling if their lipid species to adjust to growth temperature change^24^, when cultivated on undefined lipid diet. Fig.5a shows that mycoplasma maintains its ability to engage in active lipidomic remodeling regardless lipidome size and composition. Furthermore, the magnitude of total lipidomic remodeling of all lipid species in both *M. mycoides* and Syn3B (with the only exception of 4FAs-grown Syn3B cells) remains relatively conserved, in 10-25 mol% range – comparable to cells, grown on FBS^24^. What lipidomic features underlie temperature adaptation in *M. mycoides* and Syn3B? We addressed this question by dissecting adaptive lipidome remodeling for each of the lipid diets (see data analysis methods section for how mol% remodeling was calculated) by lipid class and acyl chain features (Fig.5b, c). By aggregating the lipidome according to class we can resolve how cholesterol varies relative to phospholipids, and how phospholipid head groups are regulated independent of acyl chain composition. In *M. mycoides*, class variability is relatively conserved in all lipid diets and does not exceed 5 mol% for both cholesterol (Fig.5b) and phospholipids (Fig.5c, top). Although this pattern of class remodeling is generally similar in Syn3B, cholesterol variability tends to be slightly higher (with up to 10 mol% in cells, grown on 4FAs diet – Fig.5b), with ∼9 mol% change in PG class (2FAs diet) and CL class (4FAs diet) (Fig. 5c, bottom). Slightly higher class variability, especially cholesterol, might be an alternative mechanism for temperature adaptation in the minimal cell, which features less flexibility in phospholipid synthesis and remodeling (Fig.3).

**Figure 5.**
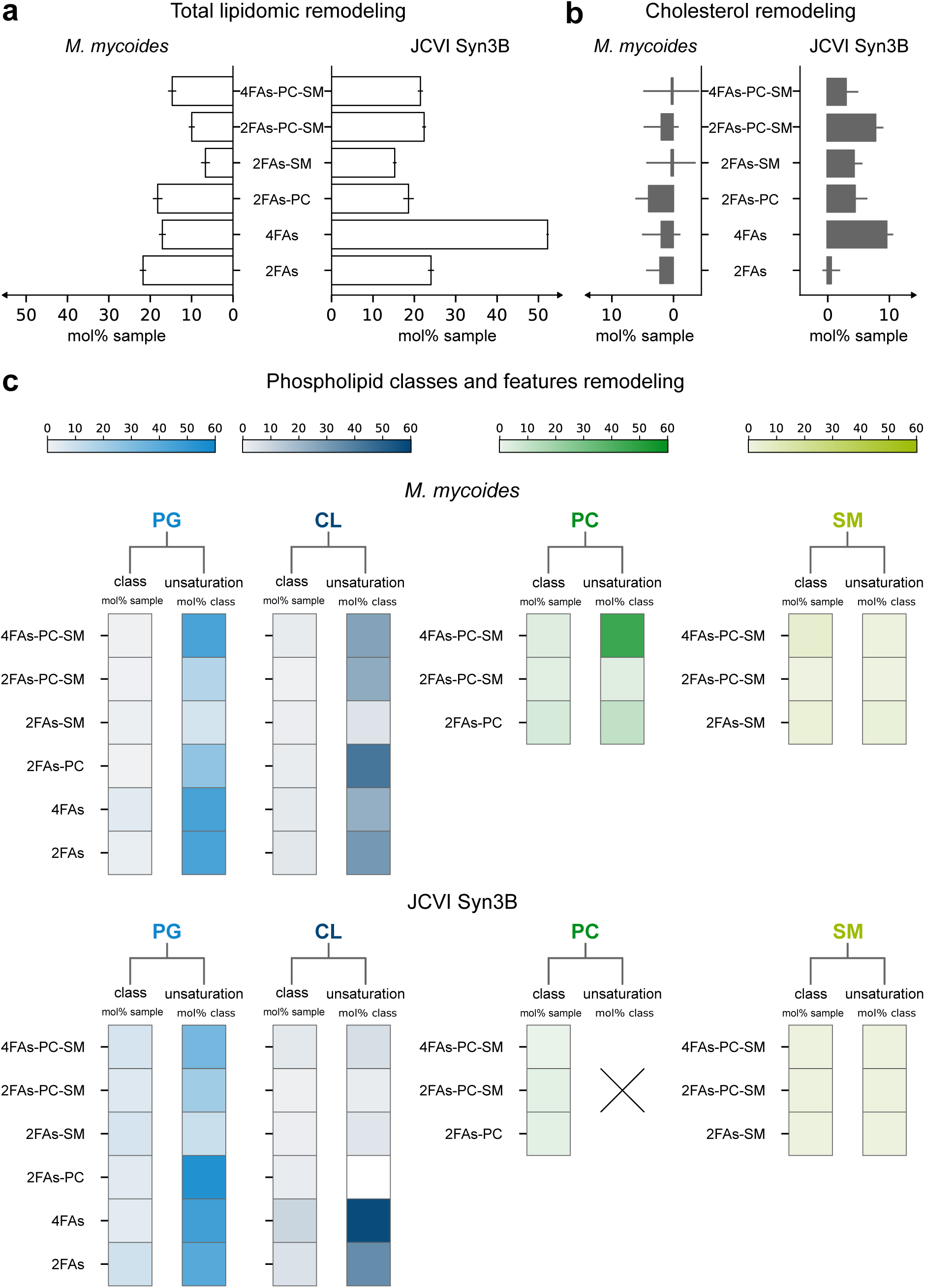
Patterns of lipidomic remodeling in mycoplasma. **a.** Total lipidomic remodeling of mycoplasma membrane lipidomes in 37°C-30°C. **b**. Cholesterol remodeling with temperature. **c**. Phospholipid classes and phospholipid unsaturation remodeling with temperature. The heatmaps show average absolute remodeling magnitude with temperature, n = 3.

#### Phospholipid acyl chain variability

We can further aggregate the lipidome by acyl chain features such as unsaturation (number of double bonds) and length (number of carbons), to reveal how acyl chains are remodeled independent of head group. Remodeling of lipid acyl chains is one of the primary means by which cells adapt their membrane properties to varying conditions like temperature^54,42^. Increasing the degree of unsaturation (average number of double bonds per lipid) or decreasing the average chain length led to an increase in fluidity of the membrane that can compensate for decreased temperature.

The combination of C16:0 and C18:1 acyl chains in 2FAs-containing diets implies a strict co-dependence of resulting phospholipid acyl chain features, making it sufficient to only consider one parameter to understand the dynamics of acyl chain features adaptation with temperature in mycoplasma, cultivated on 2FAs-based diets. The addition of C16:1 and C18:0 in 4FAs-based diets breaks this strict relationship between acyl chain length and unsaturation, allowing for larger degrees of freedom in acyl chain features remodeling. What advantages do extra acyl chains bring to mycoplasma mechanisms of lipidomic temperature adaptation?

Both *M. mycoides* Syn3B employ substantial unsaturation remodeling of internally synthesized PG on all diets, except (2FAs-SM). Cardiolipin unsaturation remodeling, however, is more divergent between two organisms and appears more lipid-diet dependent (Fig.5c). When only fatty acids are present in the diet, Syn3B actively modulates both internal phospholipid classes, and 4FAs diet results in more active remodeling of both PG and CL. When PC and/or SM are present in the diet, the minimal cells reduced cardiolipin remodeling to a minimum, with zero unsaturation change on (2FAs-PC) diet. This trend complements our observations of Syn3B producing smaller variety of CL species at 37°C (Fig.3). At the same time, the minimal cell continues to only incorporate POPC in the lipidome at 30°C, which results in the complete absence of acyl chain remodeling in this class. Remarkably, while *M. mycoides* is capable of unsaturation remodeling in PC class, it is not actively employed on 2FAs-containing diets; only 4FAs-PC-SM diet allows *M. mycoides* to engage in considerable PC acyl chain remodeling (around 50 mol% class). It might therefore be possible that the restricted diversity of acyl chains in the diet forces *M. mycoides* to rely more on internally synthesized phospholipids adaptation. The minimal cell lacks an ability to remodel the external PC entirely, as it’s only capable of incorporating the supplemented POPC. Finally, neither of two organisms perform a considerable SM acyl chain features modification at lower temperature; as the SM class composition in mycoplasma lipidomes at 37°C is identical to that of egg SM (SFig. 2b), we hypothesize that mycoplasma is incapable of acyl chain modification of sphingomyelin. Instead, the observed small changes in SM unsaturation with temperature (2-3 mol%, Fig.5c) suggest that mycoplasma chooses between available SM species in naturally derived SM. Since egg SM is heavily dominated by C16:0-SM (SFig. 2b), the rest of its species are only present in marginal quantities and did not lead to substantial SM acyl chain remodeling in *M. mycoides* and Syn3B.

Overall, Fig.5 demonstrates that *M. mycoides* and Syn3B actively remodel their lipidomes on all lipid diets, and the mechanisms of lipidomic remodeling are shaped by the constraints of lipidomic input. The presence of extra acyl chains in the diet enhances the unsaturation remodeling in both internally synthesized and externally supplemented phospholipids. At the same time, the minimal cell appears to be more restrained in phospholipid features remodeling, than *M. mycoides*, with less active cardiolipin remodeling in presence of PC and/or SM and inability of modifying PC species. Our results point at the absence of sphingolipid acyl chain remodeling in both *M. mycoides* and Syn3B. Nevertheless, substantial regulation of glycerophospholipids acyl chain features with temperature suggests the active mechanism of temperature adaptation in mycoplasma even on the minimal lipid diet. It therefore seems that chemical properties of lipid species, available to mycoplasma dictate its lipidomic remodeling patterns and adaptation. What role – if any - lipidome size plays in mycoplasma membrane properties and adaptation?

### 6. Lipidome size and composition shape mycoplasma lipidome organization and cellular fitness

We have thus far demonstrated that mycoplasma actively remodels its lipidomes at lower growth temperature on all lipid diets, and chemical properties of lipid inputs largely dictate the patterns of lipidomic remodeling and, subsequently, homeoviscous adaptation. What is the role of lipidome size in processes of lipidomic remodeling and adaptation? Is it only the chemical specificity of lipid species that shapes mycoplasma cellular fitness?

In our previous work we have revealed a universal organizational principle of membrane lipidomes, regardless their origin and specific lipid composition: in living membranes, all lipid species follow a logarithmic distribution with few species dominating the abundance space^24^. We showed that *M. mycoides* and Syn3B, despite their simplicity, follow this principle alongside tremendously more complex eukaryotic membranes. Will the logarithmic distribution of lipids hold in mycoplasma membranes, constructed on very restrained lipidomic inputs? We have sorted *M. mycoides* and Syn3B lipidomes, obtained on defined diets at 37°C and 30°C, by abundance – from the largest to the smallest- and plotted the resulting distribution against lipid species count for each lipidome. Fig.6a shows two (the largest and the smallest) lipidomes of *M. mycoides* and Syn3B, sorted from the largest to the smallest abundance (the lipidomes for the rest of lipid diets can be found on SFig. 5). Fig.6a shows that the lipidomes follow the logarithmic distribution independent from the lipid diet composition. It is remarkable that mycoplasma scales its lipids abundance logarithmically, even when only a dozen lipid species are available, which is an order of magnitude smaller lipidome, than we have recorded on animal serum^24^. However, by applying a power fit to each lipidome distribution, we have observed that the lipidome size influences the value of the power function (Fig.6b) and the respective goodness of fit (Fig.6c). The smaller lipidomes in both *M. mycoides* and Syn3B display slightly more chaotic logarithmic scaling of their lipids, with some species visibly deviating from the power line, expectedly resulting in a worse goodness-of-fit (Fig.6c). This tendency is particularly apparent in the lipidomes, approaching the lowest number of species, such as 2FAs and (2FAs-PC), suggesting that minimal lipidomes challenge the inherent membrane organization features. Interestingly, with comparable lipidome sizes, the minimal cell on average displays worse goodness-of-fit, than *M. mycoides*, which points at larger sensitivity of Syn3B to smaller number of lipid species available in the diet.

**Figure 6.**
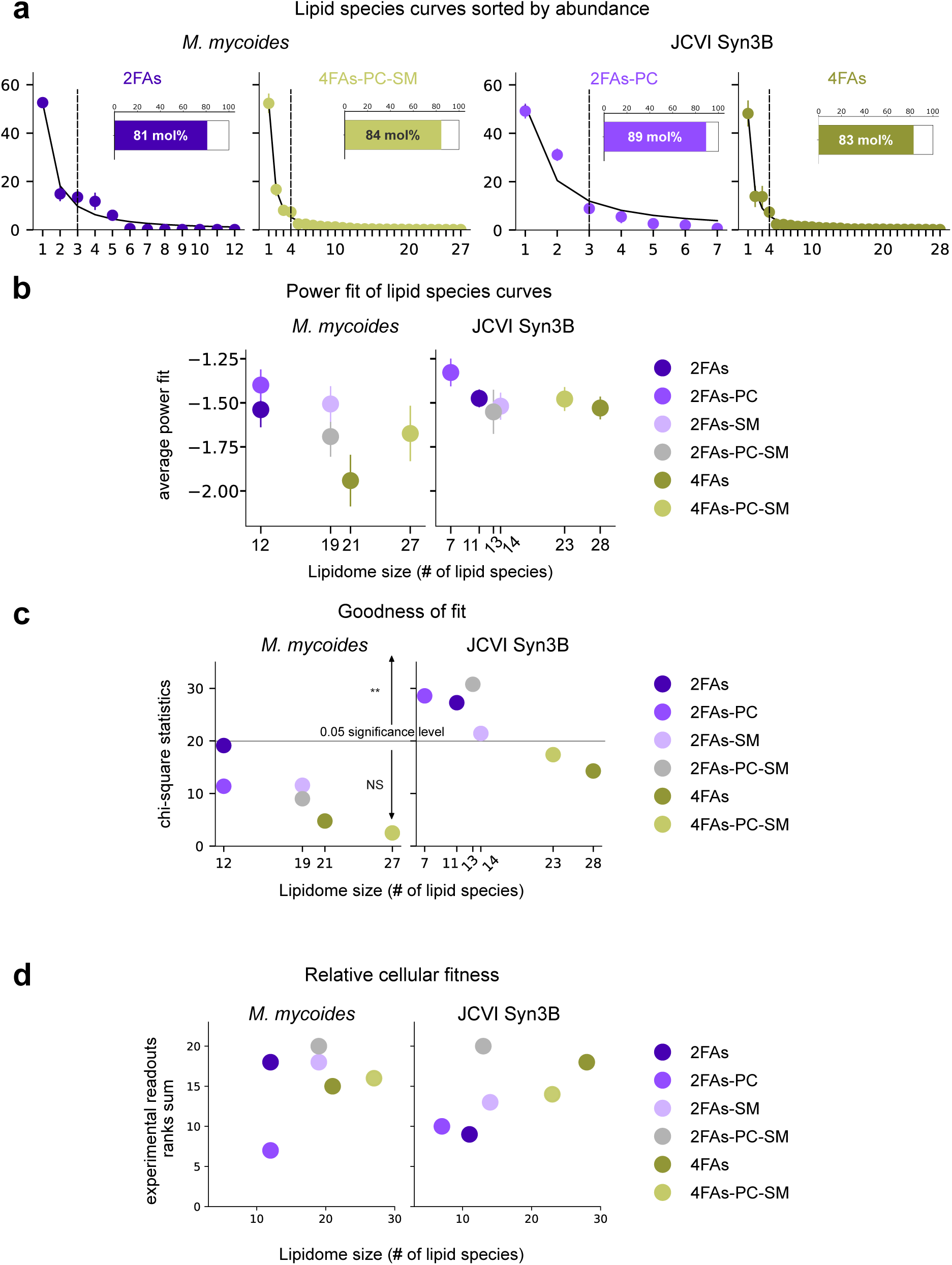
**a**. Membrane lipidome species sorted by abundance from the largest to the lowest, with an average power fit shown as a black line. Dashed vertical lines show the separation point of lipidomes into high- and low-abundance species, using Kneed algorithm (see methods for details). The inset barplots show the average abundance, covered by the high-abundance species in the sample. **b**. Numberical value for each of the samples average power fit, plotted against the respective lipidome size. **c**. Average R2 (goodness of fit) plotted against the respective lipidome size for each of the diets. For all scatter plots in **a,b** and **c** the markers represent the average value for 6 replicates (3 for 37°C and 3 for 30°C), with respective standard deviations. **d**. Relative cellular fitness, estimated as the sum of ranks, assigned to each of the experimental readouts in this study (growth rates at 37°C, homeoviscous adaptation as the difference between Pro12A GP values at 37°C and 30°C, osmotic sensitivity at 37°C and 30°C). For details see methods.

Larger lipidomes tend to produce a more negative power fit, indicating that fewer species dominate larger portion of the lipidome (Fig.6b). From our previous observations, more negative fits seem to be a common feature of cholesterol-dominated membranes^24^; here, by tuning mycoplasma membrane lipidome within an order of magnitude size range, we show that lipid species availability in the diet further steepens the logarithmic distribution, which is seemingly counterintuitive. To explain this observation, we applied the same approach to split the lipidome into two groups of high and low abundance that we have already utilized in our previous work^24^. The distribution of lipid species abundances has a single point of maximum curvature that occurs at 3-10 mol% average lipid abundance (Fig.6a). This point visually separates the lipidome into two distinct ‘tails’ of high abundance (on the left) and low abundance (on the right). The presence of the maximum curvature point indicates a transition in the abundance distribution behavior from the few dominant high-abundance lipids to the bulk of low-abundance species. To precisely locate this point on lipid distribution curves, we searched for the maximum curvature using Kneedle algorithm, originally developed by Satopää et.al^55^. The dashed line on Fig.6a shows the resulting curvature point. This line consistently separates 3-4 most abundant species from the rest of the lipidome in all the samples. This analysis reveals that considerable variability in lipidome size does not change the high abundance ‘tail’ architecture, which appears to be conserved, alongside the constant ∼50 mol% cholesterol. Consequently, high abundance tail consistently takes around 80 mol% of total lipidomic abundance, in agreement with Pareto distribution^56^ (the inset bars on Fig.6a). Therefore, when more lipids are available in the diet, both *M. mycoides* and Syn3B incorporate them in a bulk of low abundance lipidome ‘tail’, which is ultimately demonstrated by a more negative power fit. This observation points at a crucial principle of membrane lipidome composition in mycoplasma, flexible enough to withstand large fluctuations in the number of lipids available, as well as their chemical properties.

The goodness-of-fit (Fig.6c) of lipid species curves, however, is not sufficient to conclude whether lipidome size is crucial for membrane adaptation and cellular fitness. The poorer fit quality might simply be explained by the low sample number, which inevitably reduces the quality of statistical testing^57^. To examine whether lipidome size enhances the cellular fitness regardless of lipid composition, we considered all the experimental readouts in this study (growth rate at 37°C, osmotic sensitivity at 37°C and difference in Pro12A GP with temperature) to calculate a cumulative cellular fitness score. The highest rank = 6 is assigned to the best experimental readout among lipid diets, and lowest rank = 1 to the worst readout, respectively (see methods for the calculation details). The rank sum is the cumulative fitness score *M. mycoides* and JCVi-syn3B achieve on a lipid diet. The higher sum corresponds to the better cellular fitness. Fig.6d illustrates that larger lipidome size on average supports better cumulative cellular fitness in both bacterial strains. Interestingly, the minimal cell exhibits a simpler, more linear correlation between size and fitness, with the prominent exception of 2FAs-PC-SM diet. For *M. mycoides*, several diets yield identical sample sizes with very different relative fitness. These results demonstrate that regardless the lipid composition, larger lipidome sizes tend to better support mycoplasma membrane adaptation and cellular robustness, with the minimal cell being more sensitive to the reduced number of species in the membrane lipidomes. *M. mycoides* is more flexible in its lipid synthesis and remodeling patterns and is less dependent on the lipidome size; lipid composition is more substantial factor in shaping *M. mycoides* cellular fitness. The best relative cellular fitness of cells, grown with 2FAs-PC-SM diet suggests that the combination of both lipidome size and composition defines the best membrane biophysical state.

### 7. Mapping the relationship between lipid diet and lipidomic features

In this study, we explore how changing the lipid diet provided in the growth medium influences the cellular lipidomes. In this sense, lipid diets are the input, and the lipidome is the output, allowing us to characterize the lipidome remodeling capacity of *M. mycodies* and Syn3B. We’ve explored lipid remodeling in terms of lipid classes and class-specific acyl chain features. Now we take a step back and examine how the total acyl chain features vary with lipid input. The acyl chain profile alone can neither define nor explain membrane phenotypes; however, it’s a crucial parameter in defining membrane thickness^63^, curvature^64^ and lipid-protein interactions^65^. The way we have designed the lipid diets allows for varying independently the acyl chain profile derived from uptake of free fatty acids and/or acyl chains from externally delivered PC/SM (Fig. 7a and b: SFig.4a shows the average acyl chain parameters for each of the external phospholipids, while Fig.1b - for 2FAs and 4FAs inputs). On the minimal 2FAs diet at 37°C, both strains synthesize PG and CL such that their average acyl chain length and unsaturation are strictly coupled, which is visualized by values falling along a 45° line, or isocline Fig.7a and b (gray line). Both *M. mycoides* (Fig.7a(i)) and Syn3B (Fig.7b(i)) on average tend towards for shorter, more saturated acyl chains, when forced to grow on the minimal diet (square markers on the plots). Once POPC is added to the diet, both average length and unsaturation increase drastically, shifting towards the average acyl chain profile of the external lipid POPC. The prominent effect of POPC on average acyl chain features is due to our observation that both *M. mycoides* and Syn3B decrease the internal lipid synthesis in the presence of external phospholipids, incorporating up to 40 mol% of POPC or SM (Fig.3a). The average acyl chain profile of phospholipids follows (and indirectly implies) the lipid class mol% changes as a result of changes in lipid input. At the same time, 2FAs-POPC diet, as predicted, does not break the strict codependence of acyl chain features, and resulting average profile is only capable of shifting along the isocline. If, instead of POPC, we introduce additional types of free fatty acids (C16:1 and C18:0) to further diversify internally produced PG and CL acyl chain configurations, the linear codependence between length and unsaturation is eliminated as now mycoplasma can vary acyl chain length and unsaturation independently (Fig.1b). *M. mycoides* uses this additional chemical degree of freedom to synthetize phospholipids that are on average both longer and relative more unsaturated (Fig.7b(i)). The minimal cell, however, fails to change the ratio of length to unsaturation on 4FAs diet, (Fig.7c(i)), which can be attributed to its impaired lipid synthesis and remodeling, consistent with our observations in Fig.3b and Fig.5c, respectively.

**Figure 7.**
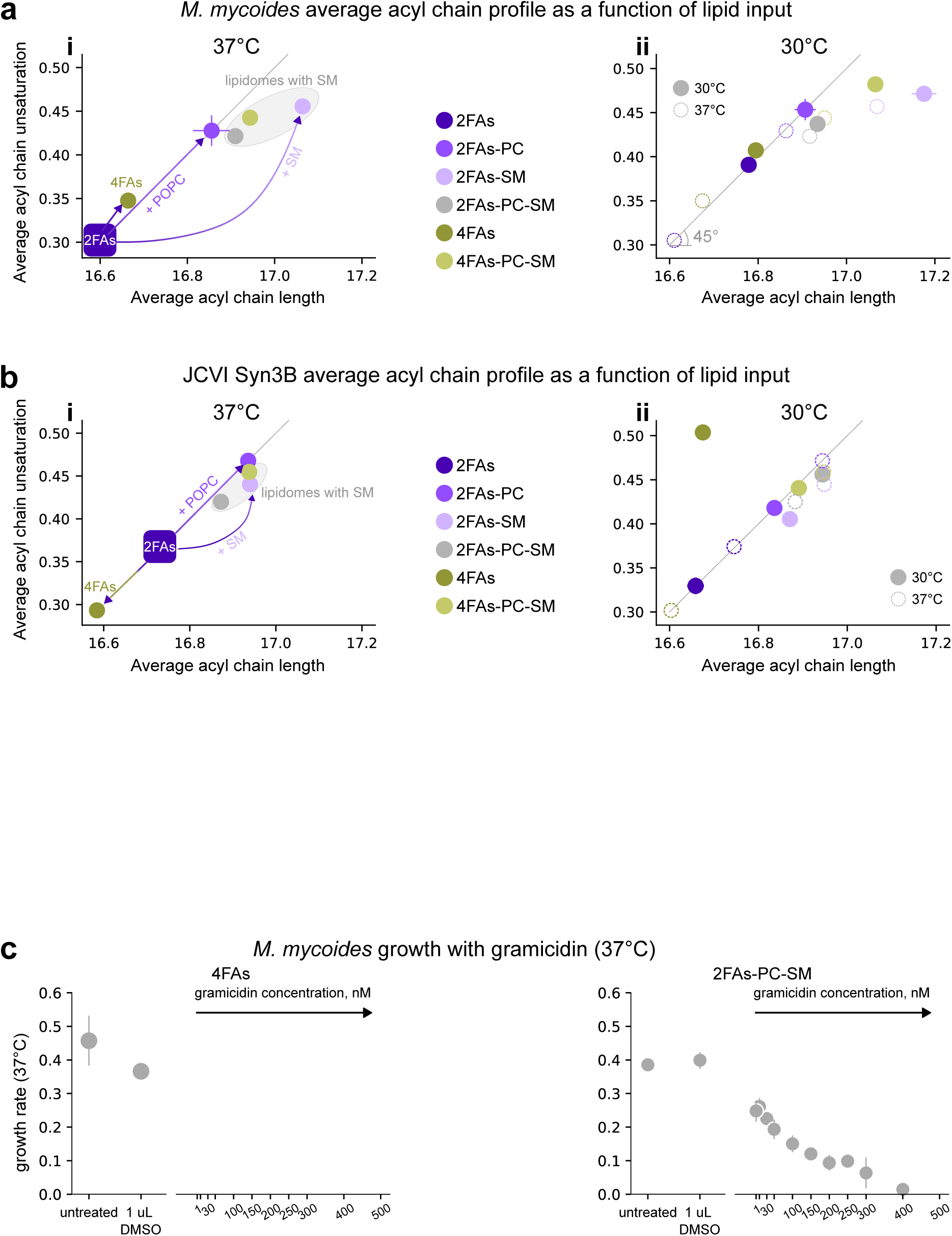
Mycoplasma is a versatile membrane model system. **a** (*M. mycoides*) and **b** (JCVI Syn3B).**i.** Average acyl chain features in the phospholipidome at 37°C (cholesterol excluded). On all subplots, 45°C line provides the reference position for features of lipidomes, grown with 2FAs. **ii**. Average acyl chain features in the phospholipidome at 30°C (solid markers). Dashed outline markers show the 37°C features, provided for comparison with 30°C.n = 3, mean +/- SD. **c**. *M. mycoides* growth at 37°C with gramicidin on two lipid diets: 4FAs (left) ad 2FAs-PC-SM (right). n = 6, mean +/- SD.

As a naturally derived sphingomyelin, egg SM used in the ‘SM’ diets contains several distinct species, shown in SFig.4a (right). As palmitoyl SM containing a C16:0 acyl chain (PSM) is the dominant species in egg SM (SFig. 2b and avanti.com), the average acyl length is only slighter longer, than in POPC, with nearly identical average unsaturation. However, it is sufficient to produce mycoplasma lipidomes that are the most deviant from acyl chains features on the minimal diet. With SM, Both *M. mycoides* and Syn3B produce the longest acyl chains, regardless the remaining lipid input composition. Since mycoplasma do not appear to be capable of remodeling acyl chains in SM (SFig. 2b), it is predictable that SM average acyl chain features will dominate the overall acyl chain profile in mycoplasma in our experimental framework.

In addition, average acyl chain analysis allows to track the changes that occur with temperature perturbation. Fig.7b(ii) and Fig.7c(ii) show the average acyl chain parameters analysis at 30°C (solid markers). 37°C points are added as dashed markers for comparison. Fig.7b(ii) shows that *M. mycoides* uses longer and more unsaturated acyl chains to adapt to lower growth temperature, regardless of the lipid input. For Syn3B, however, 30°C changes are more lipid diet-dependent, and on average less pronounced, compared to *M. mycoides*. The only striking exception is provided by the 4FAs diet: the minimal cell significantly increases average unsaturation at lower temperature. Fig.7c(ii) is, therefore, another example of how genomic minimization of Syn3B render it more sensitive to changes in lipid input in perturbations. Taken together, average acyl chain features analysis shows that it’s possible to predict a certain acyl chain profiles in the system, and demonstrates the potential to force mycoplasma to adopt predicted features by a means of a single lipid species in the diet.

### 8. Mycoplasma tunable membranes as a drug screening platform

To further illustrate the advantages of customizable model membrane lipidome composition and biophysical properties of mycoplasma membrane model system, we grew *M. mycoides* with gramicidin D (37°C). As a potent pore-forming antimicrobial peptide, gramicidin is primarily known to target gram-positive bacteria^66^. Once imbedded in the membrane, gramicidin helices form non-selective ion channels, compromising cellular cations gradients^67^. As a result, gramicidin channel conductivity is significantly higher in negatively charged lipid bilayers^68,69^. 4FAs and 2FAs-PC-SM diets yield a comparable growth rate (Fig.2a) yet a very different membrane lipid composition (Fig.3a): the only two phospholipids – PG and CL – synthesized with 4FAs diet produce a negatively charged membrane, while the addition of externally supplemented PC and SM in 2FAs-PC-SM diet produces zwitterionic bilayer. We tested mycoplasma growth with different gramicidin concentrations, ranging from 1 to 500 nM (Fig.7c). All antibiotic concentrations were added in 1uL DMSO to the final cell culture (500 uL) to exclude the additional solvent effects. Fig.7c shows the drastic differences of gramicidin on mycoplasma growth, brought upon by variations in the lipid input. As low as 1 nM gramicidin is sufficient to inhibit mycoplasma growth on 4FAs diet, while mycoplasma cells with zwitterionic membranes (2FAs-PC-SM) can withstand up to 400 nM concentration. The MIC of under 0.5 uM for 2FAs-PC-SM-grown *M. mycoides* is still considerably lower, than 2-20 uM range, utilized for other bacterial targets, such as *Staphylococcus aureus*^70^, which is likely explained by the absence of a cell wall in mycoplasma^37^. Yet variations in the lipid input in mycoplasma yield orders of magnitude larger differences in gramicidin-induced cytotoxicity. This result provides yet another evidence for the potential of *M. mycoides* tuneable membrane model system as a platform for drug testing.

## Discussion

### Mycoplasma as a tuneable membrane model system

Cellular lipidomes from microbes to mammals exhibit tremendous variations in complexity in terms of both number of lipid species and their chemical properties. Understanding the principles and conserved features underlying the diversity of membranes that have all evolved to broadly serve a similar purpose is a central challenge in membrane biology. To address this challenge, we introduce several ways to experimentally manipulate the lipidomes of minimal model organisms, providing a framework for studying the physiological and biophysical role of lipidomic features *in vivo*. We have demonstrated how defined lipid diets can be used to manipulate the lipidome composition and size in *M. mycoides* and JCVI-syn3B. The distribution of lipid classes could be adjusted by adding or removing phospholipids or sphingolipids from the lipid diet, generating lipidomes with class distributions that mimic bacterial membranes with negatively charged PG and CL, or eukaryotic plasma membranes with zwitterionic SM and PC. In turn, acyl chain diversity could be adjusted by varying the number of fatty acid species, or by introducing phospholipids with different acyl chain profile. By tuning lipid diet complexity, we could vary lipidome size from 7 to nearly 30 lipid species. The lipi omic data and experimental manipulations reported here can provide a resource for future work to understand the role of lipid complexity in minimal living membranes.

We observed that minimal cells can tolerate a nearly complete transformation in lipidome class composition, showing how a simple organism can build functional membranes from completely different lipid complements. Open ocean cyanobacteria have also been shown to transform their lipid class composition from phospholipids to glycolipids when phosphate concentrations become limiting^71,72^. These observations suggest that membrane properties essential for supporting life can be achieved through many different combinations of lipids. We can speculate that some properties can be achieved with diverse combinations of lipids, while others are only possible within a narrow range of lipid structures. For example, sterols and hopanoids are among the few lipids that can interact with phospholipids to form condensed liquid ordered membranes^73^. The unique lipid ordering properties of sterols and hopanoids are essential for many organisms to build surface membranes that are both fluid and mechanically stable^74,75^. With few exceptions, all eukaryotes have some sort of sterol, and many bacteria have hopanoids. The phospholipid class compositions of sterol- and hopanoid-containing organisms, however, can be very different^6,1^. These anecdotal observations indicate that certain properties (e.g. lipid order) can only be modulated with specific indispensable lipids, whilst other properties (e.g. fluidity) can be modulated through a broad range of lipids. Through this lens, minimal living model membrane systems offer tools to dissect the specificity of lipid structure-property relationships in different experimental framework (Fig.4 and Fig.7) that are essential to understanding the principles underlying the diversity of lipidomes across the domains of life.

### Homeoviscous adaptation

Since 1970s^42^, homeoviscous adaptation is considered an fundamental temperature adaptation mechanism in living membranes ^76,54,77,78,74^. However, its role in supporting bacterial bioactivity has recently been questioned^8,76^ and needs further investigation^79^. In our previous work, we have shown that homeoviscous adaptation is conserved, though less efficient, in the minimal cell (grown on undefined lipid diet), indicating that fluidity regulation is an essential property of living membranes^24^. By restricting the lipid input of the minimal cell, we show that fluidity regulation is not required for viability (Fig.4) as Syn3B fails to homeoviscously adapt with several lipid compositions without compromising its viability. *M. mycoides*, however, is still capable of adapting its membrane fluidity regardless the lipid diet. We postulate that homeoviscous adaptation, though not crucial at optimal growth conditions, might be important to allow cells to withstand large environmental changes. For example, in *B. subtilis*, some antimicrobial peptides^80^ or a rapid cooling cause the membrane to phase separate, threatening many critical membrane functions^76^. Remarkably, there is evidence suggesting that cells tend to maintain the melting transition temperature of their membranes around 15-20 C below their growth temperature^81,82^. For a soil bacterium, this could provide enough buffer to prevent crossing the membrane transition temperature during diurnal temperature cycles. Therefore, while in some organisms, homeoviscous adaptation is crucial for optimizing growth limiting processes, whereas in many organisms it may serve a subtler role in protecting the cell from crossing functionally critical physical states resulting from large environmental perturbations.

### Cellular growth and fitness

Varying lipid input shapes mycoplasma growth rate, membrane robustness and properties, showcasing the essential role membrane lipid composition plays in defining cellular fitness. Although both *M. mycoides* and JCVI-syn3B are flexible in withstanding large lipid diet fluctuations, the minimal cell is generally more sensitive to changes in lipid diets. Larger sensitivity to changes in lipid input forces the minimal cell to follow the growth law^28^, thus making Syn3B a remarkable model system for exploring the dynamics between nutrients, cell size and growth. Pelletier et. al^83^ defined the foundation for identifying the membrane protein components, responsible for cell size and shape in JCVI-syn3.0. The original minimal cell – JCVI-syn3.0 – features distinct pleomorphism, with irregular cell shape and aberrant membrane-bound vesicles^84^. Pelletier et.al^83^ utilized the reverse genetics approach and restored the normal spherical morphology of the minimal cell by adding 19 genes from the parent organism (*M. mycoides*). The resulting version of the minimal cell, JCVI-syn3A/B contains 7 genes, essential for spherical cell shape and conserved cell size, with at least 4 genes coding for membrane-associated protein components^83^. By turning *M. mycoides* and Syn3B into a membrane model system with tunable lipidome, we provide a complimentary tool to explore lipid-protein interactions that control cellular size and shape.

Why are lipidomes so large? The diversity of lipid structures found in cellular membranes provides a chemical space that can accommodate selectivity for signaling and membrane identity, allosteric regulation of membrane proteins, as well as means to tune a diverse array of membrane physical properties and cellular fitness. Together with growth rate, cell size control is an integral part of cellular fitness^85^. It’s encompassed by a vast number of metabolic parameters, making it challenging to assess experimentally. As a result, the growth rate is often considered as a proxy for cellular fitness^86^, and the link between membrane lipidome and the growth rate is the best one studied up to date^8,76^. Structure-property-function relationships have been studied for individual lipids through in vivo genetic manipulation or in vitro reconstitutions. These studies revealed the crucial role of various lipid species in shaping other aspects of cellular fitness, such as cellular cytotoxicity^87^, ER functionality^88^ and disease^89^. However, the collective role of lipidome size is experimentally challenging to study, particularly in a single living organism. The minimal model organisms we explore here provide a first step towards studying how lipidome size matters (or does not) in living cells.

To establish the link between cellular fitness and lipidome size, we utilized the principle of composite fitness score, employed in assessment of physical fitness in response to environmental and metabolic stressors^90,91^. To this end, we considered a qualitative estimate of fitness that summarizes different experimental readouts. While the chemical properties of lipid diets have a profound effect on cellular mechanical robustness and homeoviscous adaptation, we hypothesized that the lipidome size is an important factor in overall cellular fitness. *M. mycoides* is more resistant to fluctuations in the lipidome size, when only the minimal lipidome size results in a compromised cellular fitness; JCVI-syn3B exhibits a more positive correlation between lipidome size and fitness, suggesting that the number of lipid species alone may be an intrinsically important feature. Ultimately, our observations indicate that it will be necessary to understand how structure-property relationships change relative to varying lipidome size. In this regard, minimal living membrane systems such as *M. mycoides* and Syn3B will be essential for deciphering the fundamental role of lipidome complexity.

### Mycoplasma as a complementary model system for cell membrane simulations

The complexity of cell membranes poses a challenge to experimental probing of membrane biophysical state and membrane lipid-protein interactions, both in vitro and in vivo^58^. To this end, molecular dynamics (MD) simulations have emerged as a powerful tool to provide insights into lipid membrane physical properties and molecular interactions^59^. However, computational costs and force field limitations usually constrain membrane lipidome size to under very few distinct lipids in simulations^60,59^, with a single component systems widely utilized^61,62^. These constraints may encompass at least an order of magnitude difference in lipidome size between simulated membranes and their living counterparts, resulting in model oversimplification. In contrast, the limited lipid repertoire of mycoplasma membranes have potential to bridge the complexity of cell membranes with the compositional constraints of MD simulations.

Lipidomic features analysis provides one example of how mycoplasma with tuneable lipidomes can serve as a platform for MD simulations. As a versatile lens for lipidome dissection and analysis^2^, lipidomic features method reduces the complexity of cellular lipidomes to a few key structural components that define the chemical and physical properties of membranes. It can thus serve as a tool in developing a set of physical numeric parameters that define the membrane system. This approach helped to reveal the role of head group-specific acyl chain remodeling in temperature adaptation in conventionally-grown mycoplasma lipidomes^24^ and can be further utilized for a tuneable mycoplasma membrane of only 10-20 species.

### Mycoplasma as a drug screening platform

Our proof-of-principle growth experiment with gramicidin shows how manipulating membrane lipid composition in mycoplasma can be utilized to explore membrane-dependent growth inhibitory effects of cationic peptides. As bacterial resistance poses a threat to global health^92^, further exacerbated by challenges discovering new antimicrobial compounds^93^, identifying new ways of drug screening remains a fervent task. As a mammalian parasite, in its native habitat mycoplasma has a unique membrane that combines typical features of eukaryotic and prokaryotic membranes. By taking control of mycoplasma membrane composition, we can force mycoplasma lipidome features closer to mammalian plasma membrane or bacterial membranes, essentially creating different platforms for testing properties of different compounds on the same cellular chassis. The versatility of this system will not only allow for effective predictions of drug inhibitory effects but also a more direct assessment of drugs’ inhibitory mechanisms at the membrane interface.

## Acknowledgements

The authors thank members of the Sáenz lab, Tomasz Czerniak, Isaac G. Justice, and Ha Ngoc Anh Nguyen, as well as Ilya Levental and Kandice Levental for their valuable discussions and feedback. Additional thanks go to our colleagues at the J. Craig Venter Institute (JCVI)—John Glass and Kim Wise— for their feedback and manuscript comments. We are grateful to Telesis Bio, Inc., for providing the bacterial strain JCVI-syn3B, and to Lipotype GmbH for their generous assistance with lipidomic sample preparation and analysis. Finally, we thank Andrey Klymchenko and Dmytro Danylchuk from the Faculté de Pharmacie, Université de Strasbourg, for providing Pro12A for lipid order measurements. This work was supported by B CUBE at TU Dresden, a grant from the German Federal Ministry of Education and Research (BMBF) (to J.P.S., project 03Z22EN12), and a VW Foundation Life grant (to J.P.S., project 93090).

## Methods

### Bacterial strains

*M. mycoides subsp. capri strain GM 12*

JCVI-syn3B

Both bacterial strains were received from J. Craig Venter Institute (JCVI) (La Jolla, California, USA)

### Chemicals

#### a. Bacterial growth medium

Both mycoplasma strains were cultivated in SP4 growth medium, supplemented with CMRL. The original SP4 recipe was first developed by Tully et.al in 1970s ^62^; later, colleagues from J.Craig Venter Institute supplemented SP4 with extra nutrients to support viability of the minimal cell^95^. The detailed SP4 composition is listed in Table 1 in Supplementary Materials.

#### b. Cyclodextrins

mßCD was obtained from Sigma. Stored at 4°C.

mαCD was purchased from CycloLab. Stored at 4°C.

80 mM stocks of each cyclodextrin were dissolved in PBS, pH-ed to 7.0 and stored at 4°C.

#### c. Lipids

Cholesterol (Sigma)

Palmitic acid (C16:0, Sigma)

Oleic acid (C18:1, Sigma)

Palmitoleic acid (C16:1, Sigma)

Stearic acid (C18:0, Merck)

POPC (Avanti)

Egg SM (Avanti)

All lipids were purchased in pure form and stored at -20°C prior use. To make working lipid stocks:

1. POPC and egg SM were dissolved in pure EtOH to 100 mg/ml concentration. The precise mM concentration was further confirmed with phosphate assay.
2. Cholesterol and fatty acids were dissolved in chloroform to yield concentrated stocks of the following concentrations:

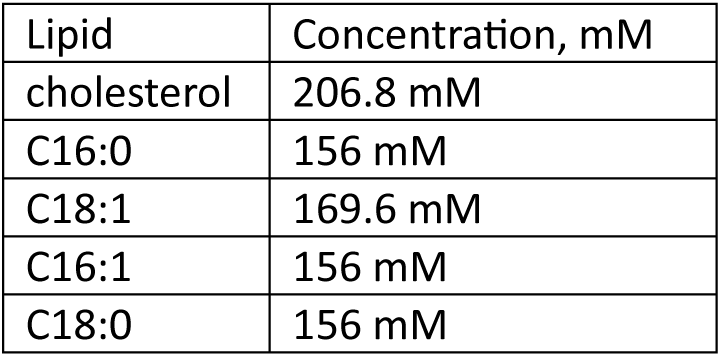

The chloroform stocks were kept in sealed glass vials at -20°C.

1. Combined working EtOH stocks with cholesterol and fatty acids were prepared as follows:

a. Total lipid concentration was 211.1 mM:

|  |  |  |  |
| --- | --- | --- | --- |
| 2FAs stock | Concentration, mM | 4FAs stock | Concentration, mM |
| cholesterol | 51.7 mM | cholesterol | 51.7 mM |
| C16:0 | 78 mM | C16:0 | 39 mM |
| C18:1 | 81.4 mM | C18:1 | 42.4 mM |
|  |  | C16:1 | 39 mM |
|  |  | C18:0 | 39 mM |

Lipids in chloroform were combined in 2 ml solvent-proof Eppendorf tubes and the solvent was evaporated using vacuum concentrator for 40 min. Next, 1 ml of pure EtOH (Uvasol) was added to the lipid film and the lipid stocks were incubated at 50°C for 20 min to ensure complete solubility of the lipid film.

#### d. Fluorescent dyes

Pro12A dye was a courtesy of Klymchenko Lab^45^. 0.5 mM DMSO stock is stored at -20°C. Propidium iodide was purchased from Cayman Chemical in pure form, stored at 4°C.

### Cyclodextrin-lipid complexes and bacterial lipid feeding

1. During SP4 growth medium preparation, total H2O volume was reduced by 1/20 to leave space for cyclodextrin incorporation later: per 1L total SP4, 950 ml stock was prepared.
2. 2FAs and 4FAs stocks are complexed with mßCD and phospholipids – with maCD. To ensure best cyclodextrin-loading, different cyclodextrins are always incubated with their respective lipid types separately.
3. For all cyclodextrin-lipid complexes, the lipid stock in EtOH was transferred to an eppendorf tube and the solvent was dried under vacuum. Cyclodextrin stocks were heated to 37°C and added to the dry lipid film. The samples were left overnight at 40°C with 1000 rpm shaking to ensure complete dissolution of dry lipid film. The complex should be a homogenous, transparent solution. Complexed stocks were stored at -20°C and heated to 37°C prior use. The complexed stocks are to be used within a week.
4. Cyclodextrin-lipid stocks in final SP4:

*M. mycoides:*

1. mM mßCD:0.2 mM 2FAs/4FAs (for all diets)
2. mM maCD:0.04 mM POPC/egg SM - for 2FAs-PC and 2FAs-SM diets
3. mM maCD:(0.02 mM POPC + 0.02 mM egg SM) - for (2FAs/4FAs-PC-SM) diets

JCVI-syn3B:

1. mM mßCD:0.2 mM 2FAs/4FAs (for all diets)
2. mM maCD: 0.025 mM POPC/egg SM - for 2FAs-PC and 2FAs-SM diets
3. mM maCD: (0.0125 mM POPC + 0.0125 mM egg SM) - for (2FAs/4FAs-PC-SM) diets

5. Cyclodextrin-lipid complexes were prepared at 40x concentration to be diluted in SP4 medium. For final 10 mL of SP4 culture the recipe is as follows:

| For 2FAs/4FAs diets (no phospholipids) |  |  |  |
| --- | --- | --- | --- |
| Final SP4 volume with lipids, ml | Add SP4 stock, ml | Add mβCD:FAs | Add PBS |
| 10 | 9.5 | 250 μL | 250 μL |
| For diet with FAs and phospholipids |  |  |  |
| Final SP4 volume with lipids, ml | Add SP4 stock, ml | Add mβCD:FAs | Add maCD:PLs |
| 10 | 9.5 | 250 uL | 250 uL |

1. To prevent lipid precipitation in the growth medium, SP4 was warmed to 37°C prior adding cyclodextrin-lipid stocks.

#### Determination of membrane lipidome composition using shotgun lipidomics

##### 1. Whole cell sample preparation for lipidomic analysis

Bacterial samples were grown in batch cultures (in triplicates) at 37°C and 30°C and harvested at early-to-mid exponential growth stage (*M. mycodies*: OD 600 0.2-0.3, Syn3B: OD 600 0.07-0.13) and washed twice in Mycoplasma wash buffer (200 mM NaCl, 25 mM HEPES, 1% glucose, pH 7.0) to remove traces of the growth medium at their growth temperature (21000g, 2 min centrifuging steps). A respective blank with SP4 + lipids was incubated together with bacterial cultures, washed and processed in the same way as the bacterial samples. Washed cell pellets and blank samples were resuspended in 150 uL PBS and flash-frozen in liquid nitrogen.

##### 2. Lipid extraction for mass spectrometry lipidomics

Lipids were extracted using a two-step chloroform/methanol procedure^96^. Samples were spiked with internal lipid standard mixture containing: cardiolipin 14:0/14:0/14:0/14:0 (CL), ceramide 18:1;2/17:0 (Cer), diacylglycerol 17:0/17:0 (DAG), hexosylceramide 18:1;2/12:0 (HexCer), lyso-phosphatidate 17:0 (LPA), lyso-phosphatidylcholine 12:0 (LPC), lyso-phosphatidylethanolamine 17:1 (LPE), lyso- phosphatidylglycerol 17:1 (LPG), lyso-phosphatidylinositol 17:1 (LPI), lyso-phosphatidylserine 17:1 (LPS), phosphatidate 17:0/17:0 (PA), phosphatidylcholine 17:0/17:0 (PC), phosphatidylethanolamine 17:0/17:0 (PE), phosphatidylglycerol 17:0/17:0 (PG), phosphatidylinositol 16:0/16:0 (PI), phosphatidylserine 17:0/17:0 (PS), cholesterol ester 20:0 (CE), sphingomyelin 18:1;2/12:0;0 (SM), triacylglycerol 17:0/17:0/17:0 (TAG) and cholesterol D6 (Chol). After extraction, the organic phase was transferred to an infusion plate and dried in a speed vacuum concentrator. 1st step dry extract was re-suspended in 7.5 mM ammonium acetate in chloroform/methanol/propanol (1:2:4, V:V:V) and 2nd step dry extract in 33% ethanol solution of methylamine in chloroform/methanol (0.003:5:1; V:V:V). All liquid handling steps were performed using Hamilton Robotics STARlet robotic platform with the Anti Droplet Control feature for organic solvents pipetting.

##### 3. MS data acquisition

Samples were analyzed by direct infusion on a QExactive mass spectrometer (Thermo Scientific) equipped with a TriVersa NanoMate ion source (Advion Biosciences). Samples were analyzed in both positive and negative ion modes with a resolution of Rm/z=200=280000 for MS (species structural resolution; see Table 2 in Supplementary Materials) and Rm/z=200=17500 for MSMS (subspecies structural resolution; see Table 2) experiments, in a single acquisition. MSMS was triggered by an inclusion list encompassing corresponding MS mass ranges scanned in 1 Da increments ^97^. Both MS and MSMS data were combined to monitor CE, DAG and TAG ions as ammonium adducts; PC, PC O-, as acetate adducts; and CL, PA, PE, PE O-, PG, PI and PS as deprotonated anions. MS only was used to monitor LPA, LPE, LPE O-, LPI and LPS as deprotonated anions; Cer, HexCer, SM, LPC and LPC O- as acetate adducts and cholesterol as ammonium adduct of an acetylated derivative ^98^.

Lipid extraction and MS data acquisition have been performed by Lipotype GmbH.

##### 4. Lipidomic data filtering

Each sample’s data was obtained in 3 biological replicates and each lipid species is reported as mol% of the total sample for each temperature (i.e, all lipid species sum up to 100% for each replicate).

To filter the data, we applied the cutoff based on the total average and lipidomic remodeling of lipid species between two growth temperatures (37°C and 30°C):

1. Calculate total average of each lipid species in the lipidome across both temperatures (3 replicates each temperature) and sort the average from the most to the least abundant lipid species
2. Calculate lipidomic remodeling for each lipid species with temperature according to the formula:

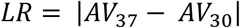

3. Sum of lipidomic remodeling for all lipid species in unfiltered sample represents total unfiltered lipidomic remodeling
4. By applying a mol% abundance cutoff from 0.01 to 100% range to the average lipidomic abundance, track how much total unfiltered lipidomic remodeling is reduced as a result. Supplementary Figure 1a shows the remodeling percentage of cutoff lipids in the total unfiltered lipidomic remodeling for both *M. mycoides* (left) and JCVI-syn3B (right). The small inset plot for each of the strains zooms into 0-0.5 mol% lipidomic cutoff.

To choose the optimal filtering abundance cutoff, we aimed to minimize the total lipidomic remodeling loss, attributed to the removed lipids. Supplementary Figure 1a shows that for both *M. mycoides* and Syn3B, 0.1-0.3 mol% abundance cutoff maintains at least 85-90 mol% of total lipidomic remodeling, which indicates that the most variable lipid species are retained in the lipidomic sample. We chose 0.1 mol% average lipid species abundance as the filtering threshold for all lipidomic samples in this study.

#### Mycoplasma growth using calorimetry

Bacterial cells were adapted to grow with each lipid diet for at least 3 passages prior the experiment (5 passages for Syn3B, cultivated on 2FAs and 4FAs diets). Cells were passaged daily to exclude ageing effects on growth. Batch bacterial cultures were setup in glass 100 ml flasks; cells were harvested at mid-exponential phase and diluted in their native growth medium (SP4 + cyclodextrin-lipid complexes) to the starting OD 600 nm = 0.005 (JCVI-syn3B) and OD600 = 0.001 (*M. mycoides*). Next, cells were transferred to calorimetry glass tubes, sealed manually with disposable tin lids. For *M. mycoides*, 250 uL of cell suspension was used, for Syn3B – 2 ml. To carry out heat flow recordings, the samples were submitted to the lab of Professor Karim Fahmy (HDZR, Dresden). The resulting heat flow curves were normalized for the final flat line values. The maximum heat flow value was used as a proxy for the maximum cell density. The log-linear slope of the exponential part of the heat flow curves was used as a growth rate estimate for *M. mycoides* and JCVI-syn3B. Growth experiments were carried out in biological triplicates.

#### Mycoplasma cell size determination using dynamic light scattering

*M. mycoides* and Syn3B cells were grown to mid-exponential stage at 37°C with all lipid diets in biological triplicates in batch cultures. 7 ml of cell culture was harvested and centrifuged to remove growth SP4 (9000g, 2 min). Cells were resuspended in 1mL of preheated MWB (200 mM NaCl, 25 mM HEPES, 1% glucose, pH 7.0) and cell suspension absorbance was determined at OD600 nm using DeNovix DS-11 FX+ spectrophotometer. Cells were diluted to OD 0.1 in MWB and incubated at 37°c for 5 min, 500 rpm shaking. Cell suspension was transferred to 1mL disposable cuvette and its precise concentration (OD 600) was again measured in DeNovix DS-11 FX+ spectrophotometer. The average cell diameter of cells in MWB was determined by Malvern Zetasizer, with the following settings:

|  |  |
| --- | --- |
| Material | Liposomes |
| RI (material) | 1.45 |
| Absorption | 0.001 |
| Dispersant | Water |
| Temperature | 37.0°C |
| Viscosity | 0.6864 cP |
| RI (dispersant) | 1.33 |

#### Membrane lipid order determination using Pro12A probe

Batch bacterial cultures in 100 ml glass flasks are grown in biological triplicates at each of the growth temperatures. Cells are harvested at early exponential growth state, (OD 600 nm 0.2-0.29 for *M. mycoides* and 0.05-0.15 for JCVI-syn3B), washed twice (12000g, 2 min) and resuspended to final OD 0.2 in mycoplasma wash buffer (200 mM NaCl, 25 mM HEPES, 1% glucose, pH 7.0). Working stock of Pro12A dye is prepared at 160 uM concentration in DMSO. 0.5 uL/200 uL cell culture was added to each well of 96-well plates to yield final 0.4 uM dye concentration. Washed bacterial cells in mycoplasma buffer are added on top of the dye to each well. The measurement plate with labelled bacterial samples is incubated at a growth temperature for 20 min, fluorescence recorded at 440 nm and 490 nm, with 20 nm bandwidth, following excitation at 360 nm +/- 10 nm. Pro12A general polarization was calculated according to the formula:

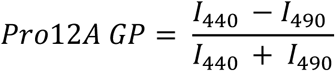

Measurements are carried out in Tecan Spark 20 multimode microplate reader, equipped with a thermostat capable of maintaining the temperature with the accuracy of +/- 1°C.

To ensure measurement reproducibility, restraining the harvesting window of cellular growth state is crucial. All consumables, wash buffer, Tecan Spark and the centrifuge are preheated to the bacterial growth temperature for at least 1 hour prior the measurement. *M. mycoides* cells are passaged twice a day. JCVI-syn3B cells are passaged daily. Cells were adapted to growth at either temperature and lipid diet for at least 3 passages (5 passages for Syn3B cells with 2FAs and 4FAs diets). After washing steps, cellular density is estimated in the buffer to yield the final density at OD 0.2 to ensure constant dye/cell density ratio.

#### Liposome preparation for Pro12A assay

Phospholipid concentration is determined prior the liposome preparation with phosphate assay. The following lipid mixtures is prepared, with total concentration of 5 mM:

1. Cholesterol:POPC 1:1
2. Cholesterol:DPPC 1:1

Lipids were mixed in solvent-proof 2 ml Eppendorf tube and chloroform is evaporated in vacuum concentrator (Christ RVC 2-25 CDplus). Dry lipid films are rehydrated in mycoplasma wash buffer (200 mM NaCl, 25 mM HEPES, 1% glucose, pH 7.0) and incubated at 55°C for 30 min, 1000 rpm shaking. Lipid mixtures are then sonicated in the sonic bath for 5 min. Finally, 10 freeze-thaw cycles are performed on all mixtures and liposomes were stored at-20°C. Liposomes are prepared in triplicates.

For Pro12A staining the same protocol is used as for mycoplasma cells. Liposomes are diluted in mycoplasma wash buffer to 100 uM concentration and stained with 0.25 mol% dye

#### Osmotic shock assay

*M. mycoides* and JCVI-syn3B cells are grown at 37°C and 30°C to early exponential state and washed once (12000g, 2 min) in mycoplasma wash buffer (200 mM NaCl, 25 mM HEPES 1% glucose, pH = 7). Washed cell pellet is resuspended in MWB to yield cell density of OD 600 nm = 0.2. Washed cells are diluted 20x (*M. mycoides*) or 10x (JCVI-syn3B) in MWB with NaCl concentration, ranging from 0 mM (=no salt) to 200 mM (default, isotonic). NaCl concentrations in MWB tested: 0 mM, 15 mM, 30 mM, 50 mM, 100 mM, 200 mM. The assay is performed in 96-well transparent bottom plates (Greiner Bio-One 655 090). First, 2 uM propidium iodide (1uL of 400 uM stock in PBS) is added to the wells. Next, 190 uL (for 20x dilution) or 180 uL (for 10x dilution) MWB with different NaCl concentrations is added on top of the dye. 10 uL (for 20x dilution) or 20 uL (for 10x dilution) of cell suspension at OD 0.2 is added to the dye-buffer mix and incubated for 10 min at the growth temperature. The final salt concentration in each well is calculated by taking into account 10/20 uL of cell suspension, added to wells. Propidium iodide fluorescence and cellular absorbance at 600 nm are measured in multimode plate reader Tecan Spark 20 at the growth temperature. Fluorescent signal is recorded at 610 nm +/- 20 nm, followed by excitation at 515 +/- 20 nm. For osmotic sensitivity data analysis, propidium iodide RFUs are normalized for the respective cellular absorbance and plotted against the salt concentration in each sample. Log-linear fit of the salt-fluorescence curves yields osmotic sensitivity estimates as the linear fit slope magnitude. The assay is performed in biological triplicates, with 3 analytical measurements per each replicate. The mean of analytical measurements is considered for sensitivity calculation. The consumables, wash buffers, centrifuges and Tecan are preheated to growth temperatures.

#### Lipidomic data analysis

*Data grouping.* For various purposes of lipidomic analysis, the filtered list of lipidomic species is grouped by different structural properties of lipids: structural category, head group, length or unsaturation, in most cases yielding the smaller subset of lipidomic entries. Therefore, the resulting subset is resummarized to 100 mol% prior subsequent analysis.

*Lipidomic remodeling.* Unless stated otherwise, lipidomic remodeling of a given lipidomic subset is a sum of absolute changes of species/features averages across 2 growth temperatures, as indicated by the following formula:

Where:

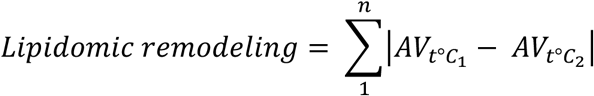

AV = species average abundance at a given temperature, n = total number of entries in a given lipidomic subset.

Total lipidomic remodeling refers to the remodeling magnitude of all species in a lipidomic sample across two different growth temperatures. Standard deviations for lipidomic remodeling are propagated using pooled standard deviation.

*Fig.6a.* Maximum curvature of lipid abundance curves is determined by kneed python package (method published by Satopaa et.al^55^)

*Fig.7, average features calculation.* Average chain length and unsaturation for each phospholipid are determined from species features according to the acyl chain distribution. The calculation is carried out according to the following examples:

PG with total length of 34 carbons and 1 double bond - PG 34:1

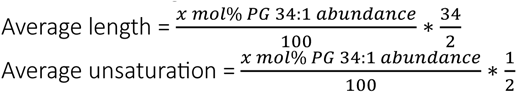

This calculation is applied to all phospholipids with two acyl chains: PG, PC and SM

CL with length of 68 carbons and 2 double bonds = CL 68:2

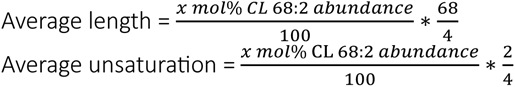

