## Supplementary Figures for "Chemically defined lipid diets reveal the versatility of lipidome remodeling in genomically minimal cells"

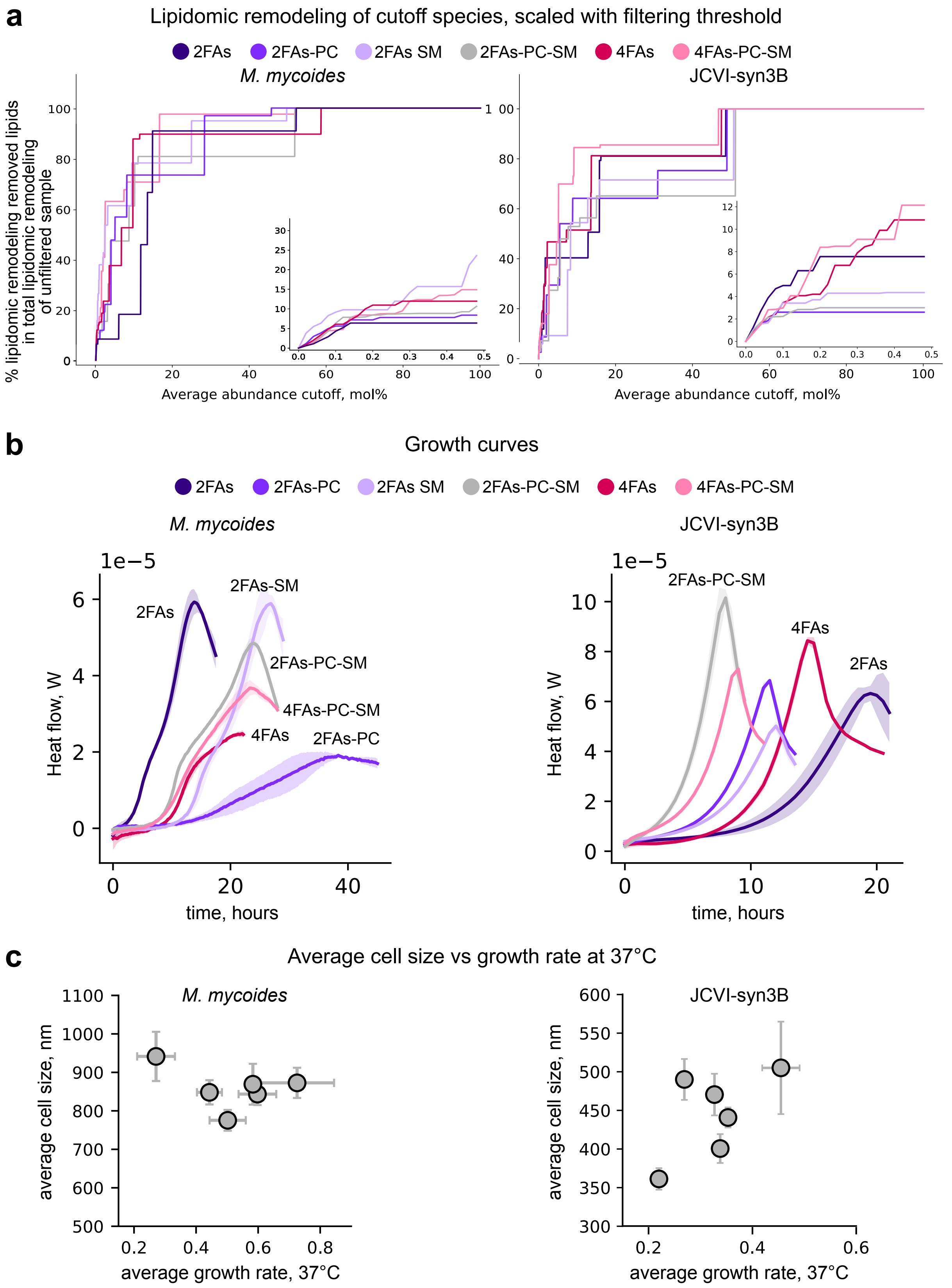


**Supplementary Figure 1. a**. Lipidomics filtering threshold in this study was decided based on % of filtered out species remodeling in total lipidomic remodeling of unfiltered lipidome with temperature for each diet. The remodeling % is plotted agaisnt average abundance filtering cutoff, mol% sample. Average was calculated for both temperatures - 37°C and 30°C, n = 6 (n = 3 for each temperature). The inset plots zoom in on the lowest cutoff mol%, up to 0.5 mol% sample. **b**. Heat flow curves of *M. mycoides* and Syn3B cells, grown at 37°C on defined lipid diets and used for growth rate calculation (**Fig.2a**). mean +/- SD, SD is shown as bands with the heat curves. **c**. Average cell size (diameter, nm) shown against average growth rate at 37°C. n=3, mean +/- SD.

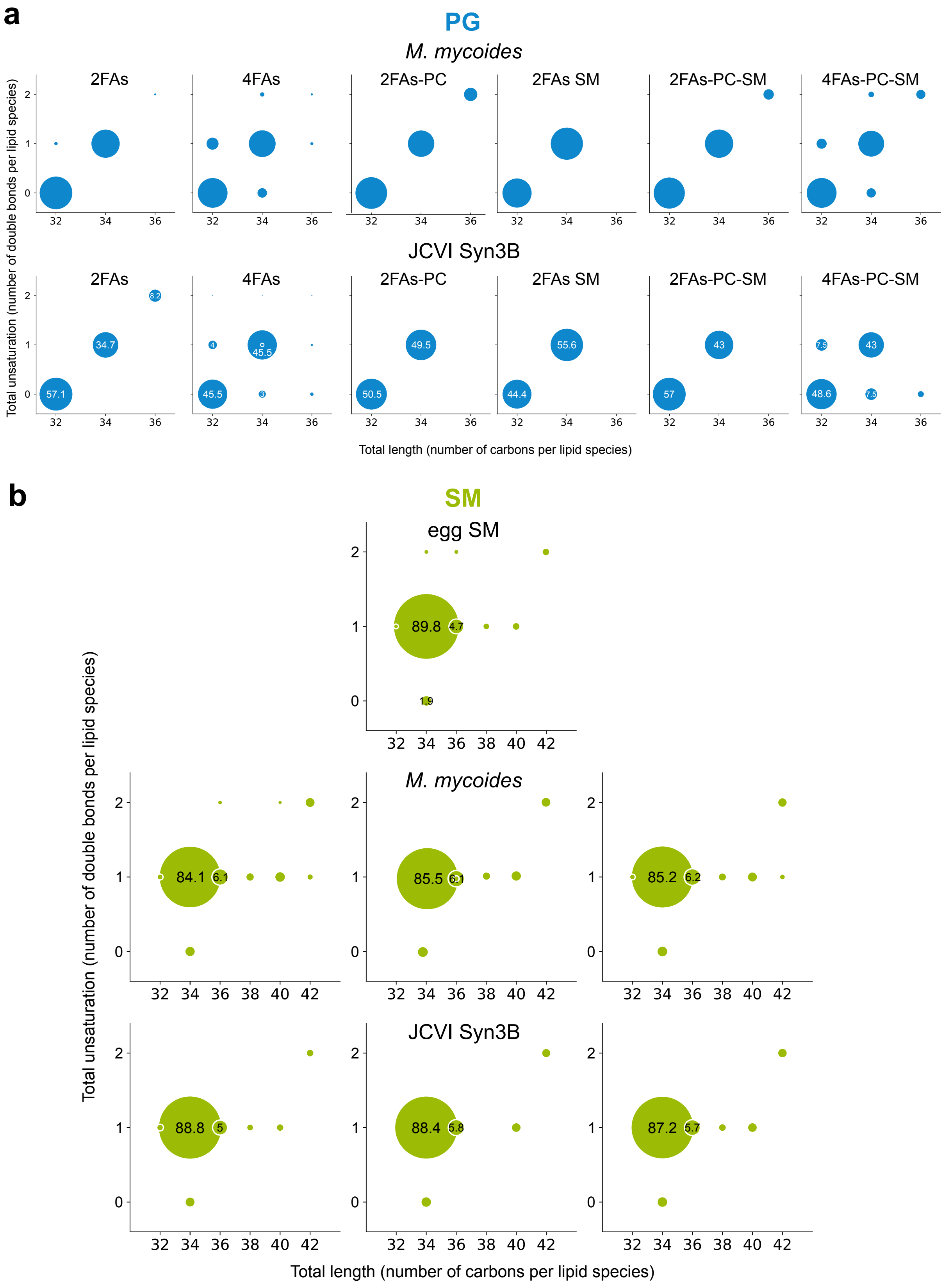


**Supplementary Figure 2. a**. PG species in *M. mycoides* and Syn3B lipidomes at 37°C. Circular plots show average abundance (n = 3), with circular diameter scaled accordingly. b. SM species distribution in egg SM (n = 1) in *M. mycoides* and Syn3B lipidomes at 37°C. Circular plots show average abundance (n = 3), with circular diameter scaled accordingly.

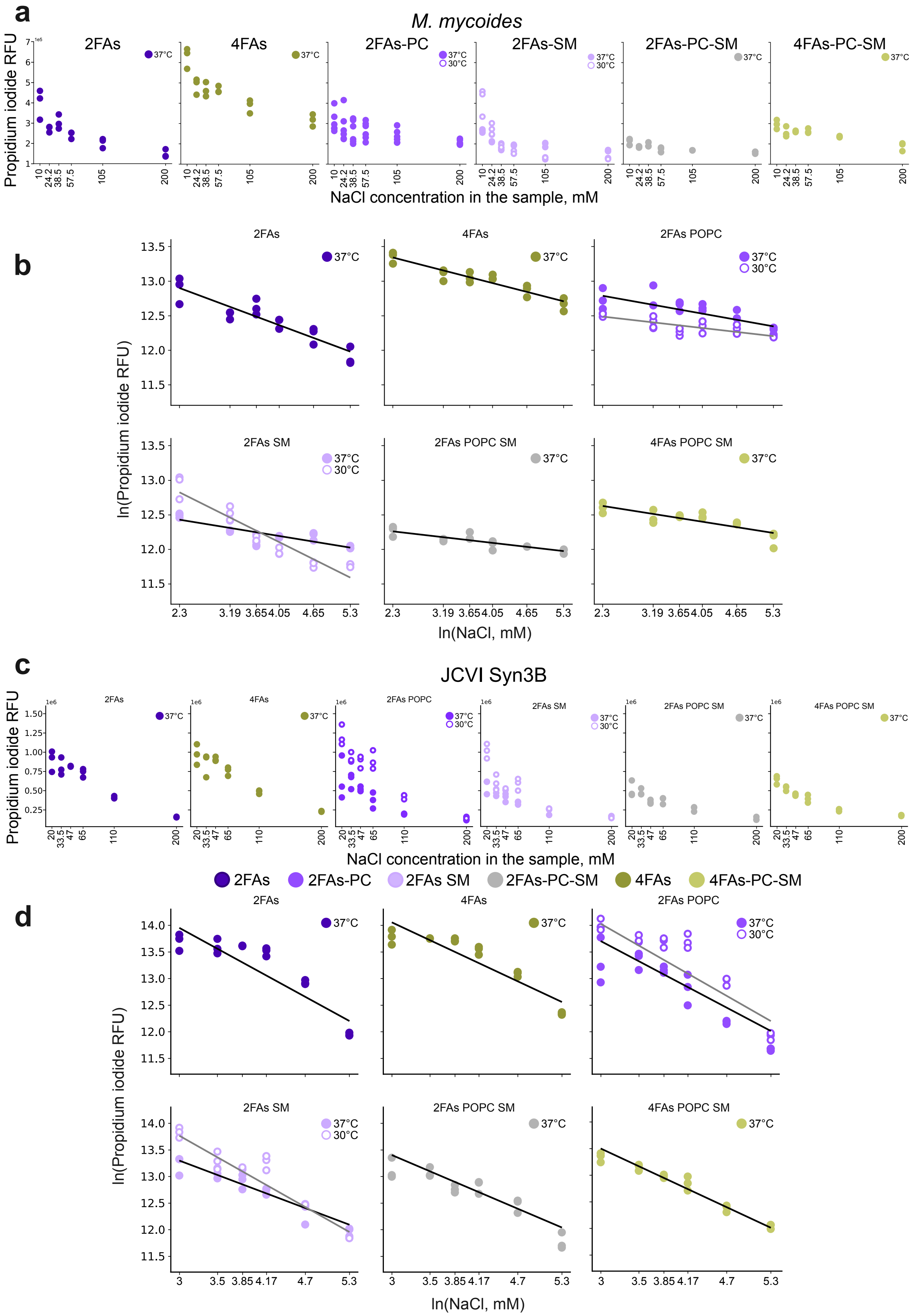


**Supplementary Figure 3.** Osmotic sensitivity assay, using propidium iodide fluorescence. **a,c**. Raw PI RFU of mycoplasma cells, grown with different lipid diets at 37°C (filled circles) and 30°C (outlined circles). n = 3. **b,d.** log values of PI RFU poltted against ln(NaCl) concetrantion in the sample with linear fit for 37°C (black line) and 30°C (gray line) readouts.n = 3

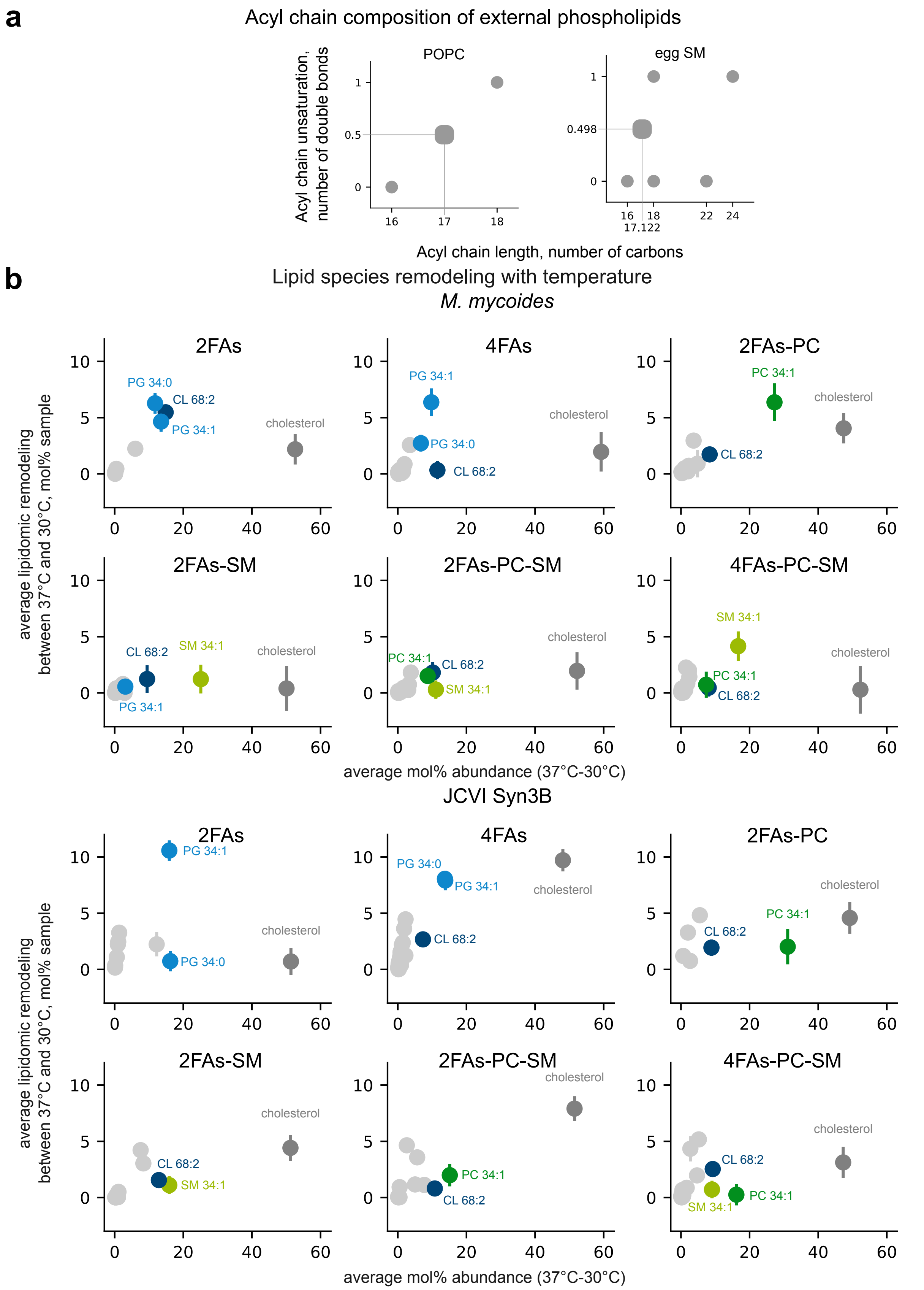


**Supplementary Figure 4. a.** Acyl chain features of all lipid inputs used in this study. Left: POPC acyl chain composition, with square marker indicating the average features in this class. Right: egg SM major acyl chain composition (data available at Avanti.com), with a square marker showing average length and unsaturation in SM class. **b.** Average lipidomic remodeling of all lipid species plotted against average abundance across both growth temperatures (37°C and 30°C). Most remodeled lipid species in each diet are color coded. n = 6 (n = 3 for each growth temperature). mean +/- SD.

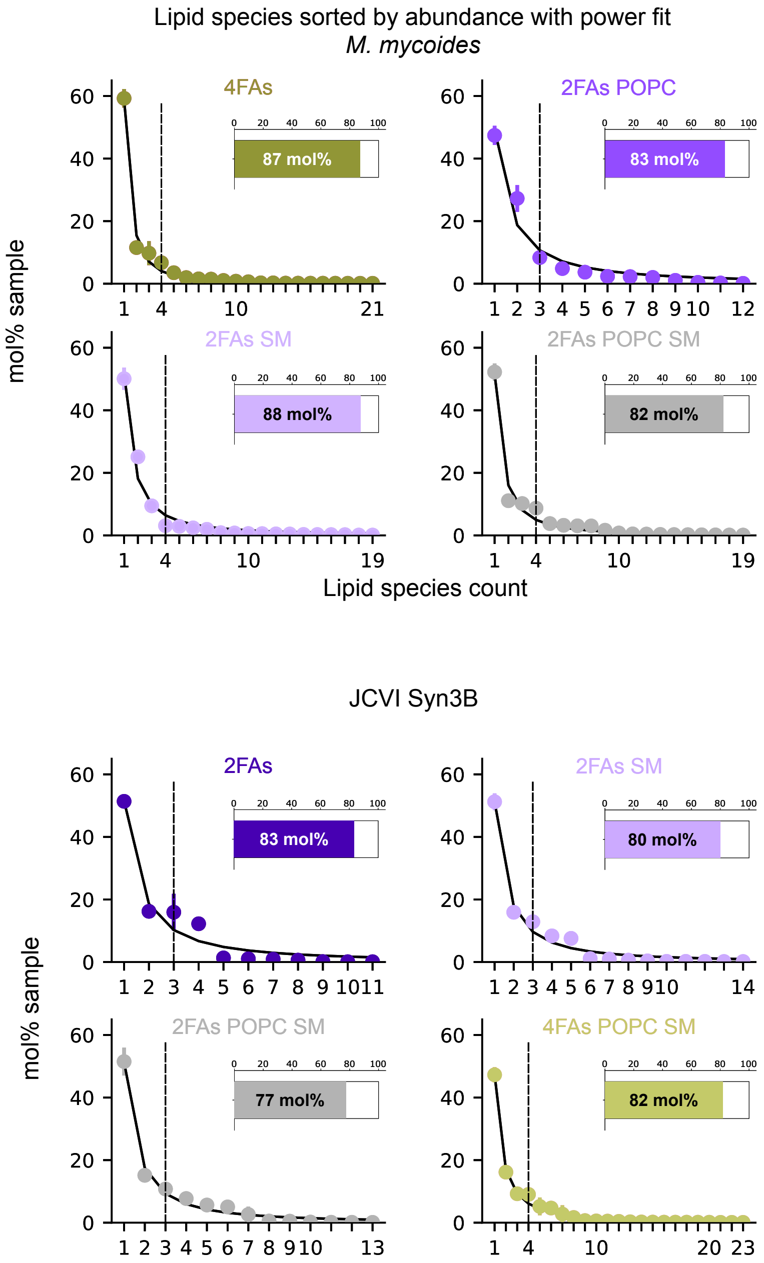


**Supplementary Figure 5**. Lipid species sorted by abundance from the most to the least abundant in *M. mycoides* (top) and JCVI-syn3B (bottom) lipidomes, grown at different lipid diets.mean +/-SD, n = 3 (biological replicates) for each growth temperature.The power fit for the average species abundance at 37°C and 30°C is shown as a black line. The rest of the diets is shown on Fig.6a.
