## Supplementary Table 1 for "Chemically defined lipid diets reveal the versatility of lipidome remodeling in genomically minimal cells"

Table 1. SP4 growth medium composition (per 1L). This medium recipe is used to cultivate *M. mycoides* and JCVI-syn3B in this study

| **SP4 growth medium, pH 7.0** | | |
| --- | --- | --- |
| Component | Amount | Company/Details |
| Difco PPLO Broth | 3.5 g | Difco/Per 1L:  Beef heart infusion from 50 g – 6 g  Peptone – 10 g  NaCl – 5 g |
| Tryptone (peptone from casein) | 10 g | Sigma Aldrich |
| Peptone (peptone from gelatin) | 5.3 g | Sigma Aldrich |
| 20% yeastolate | 10 ml | Gibco/Bacto TC yeastolate |
| 15% yeast extract | 35 ml | Carl Roth |
| 7.5% NaHCO_3_ | 14.6 ml | Honeywell |
| 20% glucose | 25 ml | Carl Roth |
| 400000 U/ml penicillin G, Na-salt | 2.5 ml | Carl Roth/Cellpure > 1550 U/mg, Penicillin G Na-salt |
| 25 mg/ml L-glutamine | 5 ml | Carl Roth/> 99% purity, Cellpure |
| CMRL | 3.92 g | HIMEDIA, CMRL 1066 Medium, w/o L-Glutamine, phenol red and sodium bicarbonate |
| Sodium bicarbonate | 0.88 g | Honeywell |
| Add H2O to the final volume of 950 ml |  |  |
